# AmpliconTyper – tool for analysing ONT multiplex PCR data from environmental and other mixed sources

**DOI:** 10.1101/2025.01.30.635642

**Authors:** Anton Spadar, Jaspreet Mahindroo, Catherine Troman, Michael Owusu, Yaw Adu-Sarkodie, Ellis Owusu-Dabo, Dilip Abraham, Benny Blossom, Karthikeyan Govindan, Venkata Raghava Mohan, Zoe A. Dyson, Nicholas Grassly, Kathryn E. Holt

**Affiliations:** Department of Infection Biology, Faculty of Infectious and Tropical Diseases, London School of Hygiene & Tropical Medicine, London, United Kingdom; MRC Centre for Global Infectious Disease Analysis, Department of Infectious Disease Epidemiology, School of Public Health, Imperial College London, London, United Kingdom; College of Health Sciences, Kwame Nkrumah University of Science and Technology, Kumasi, Ghana; The Wellcome Trust Research Laboratory, Christian Medical College, Vellore, India; Department of Community Health And Development, Christian Medical College, Vellore, India; Wellcome Sanger Institute, Wellcome Genome Campus, Hinxton, United Kingdom; Department of Infectious Diseases, School of Translational Medicine, Monash University, Australia

**Keywords:** multiplex PCR, environmental surveillance, amplicons, Oxford Nanopore Technologies

## Abstract

Amplicon sequencing is a popular method for understanding the diversity of bacterial communities in mixed samples as exemplified by 16S rRNA metagenome sequencing. This approach has been extended into multiplex amplicon sequencing in which multiple targets are amplified in the same polymerase chain reaction (PCR). Multiple tools exist to process the sequencing data produced via the short-read Illumina platform, but there are fewer options for long-read Oxford Nanopore Technologies (ONT) sequencing, or for processing data from environmental surveillance or other sources with many different organisms.

We have developed AmpliconTyper (v0.1.28, DOI: 10.5281/zenodo.14621928) for analysing multiplex amplicon sequencing data from environmental (e.g. wastewater) or similarly contaminated samples, generated using ONT devices. The software tool uses machine learning to classify sequencing reads into target and non-target organisms with very high specificity and sensitivity. The user can train models using public and/or user-generated data, which can subsequently be applied to analyse new data. The tool can also generate amplicon consensus sequences, as well as identify single nucleotide polymorphisms (SNPs) and report their genotype implications, such as association with lineages or antimicrobial resistance (AMR). The tool is freely available via Bioconda and GitHub (https://github.com/AntonS-bio/AmpliconTyper).

AmpliconTyper allows robust identification of target organism reads in ONT sequenced environmental samples, and can identify user-specified lineage or AMR markers.

**Impact statement:** AmpliconTyper (v0.1.28) is a software package that enables users to analyse amplicon sequences generated by targeted amplification followed by ONT sequencing, for environmental or other similarly contaminated samples. The analysis includes mapping of reads to target amplicon sequences, classification of each sequenced read as either originating from a target or non-target organism, followed by identification of user-specified SNPs and generation of an interactive report summarising the findings. The strength of AmpliconTyper lies in its ability to train a machine learning model using public data to create sequencing read classification models tailored to a user’s application. AmpliconTyper is designed specifically to work with extremely noisy data that includes a large share of off-target amplification reads such as those encountered in environmental surveillance applications.

**Data Summary:** For the purpose of designing and testing AmpliconTyper we have used two datasets. The first consisted of 69 *Salmonella enterica* serovar. Typhi (*S.* Typhi) and 10,303 other Enterobacteriaceae WGS ONT nanopore sequencing libraries (**Supp. Data 1**) from NCBI Sequence Read Archive (SRA) (1). We used these data to evaluate the performance of different classifier models and to train a model for our use-case, i.e. amplicon-based detection of *S*. Typhi from environmental surveillance samples (2).

In addition, to further evaluate the performance of AmpliconTyper in our use case, we have applied an amplicon sequencing protocol (2) to generate amplicon data for *S*. Typhi using two *S*. Typhi isolates (NCBI accessions SRR5949979 and SRR7165748) provided by Satheesh Nair (UKHSA) (3). We also used same protocol to the pooled sample of American Type Culture Collection (ATCC) consisting of *S*. Paratyphi A (ATCC 9150D), *S*. Paratyphi B (ATCC-BAA-1250D), *S*. Paratyphi C (ATCC-BAA-1715D), *Aeromonas hydrophila* (ATCC-7965D), *Klebsiella pneumoniae* (ATCC-BAA-1706D), and *Citrobacter freundii* (ATCC-8090D) chosen for their close relationship to *S*. Typhi. The test data for classification is available at https://github.com/AntonS-bio/AmpliconTyper/tree/main/test_data. Newly generated data was deposited in European Nucleotide Archive project PRJEB81565.

## Introduction

Polymerase chain reaction (PCR) is a well-established method for amplifying the segment of DNA lying between two primer sequences. The resulting amplicon can subsequently be subjected to sequencing to determine its precise DNA sequence, providing information on both the presence of, and variation within, the targeted segment. Multiplex PCR extends this approach by using multiple PCR primer pairs in the same reaction to generate multiple target amplicons. Multiplex PCR not only reduces costs of performing the assays, but also improves the sensitivity and specificity of organism detection, by allowing for simultaneous detection of multiple genetic regions from a single target organism (4, 5).

We have developed the AmpliconTyper software (v 0.1.28, available at: https://github.com/AntonS-bio/AmpliconTyper), to detect a specific organism based on amplicon sequences from environmental samples generated using Oxford Nanopore Technologies (ONT) devices, and also to identify lineage-defining and antimicrobial resistance (AMR)-linked single-nucleotide polymorphisms (SNPs) (6, 7). Our use case for development is detection and typing of *Salmonella enterica* serovar Typhi (*S.* Typhi), which extends previous *S.* Typhi qPCR based-surveillance work (4, 8) by integrating ONT sequencing of amplicons to more precisely detect the pathogen, and by expanding the number of *S.* Typhi-specific amplicons from 1 to 16 targeting 22 SNPs associated with 17 lineages and 4 AMR associated determinants (9). While benefiting from long reads, ONT sequencing currently suffers from substantially higher base-calling error rates compared to Illumina devices (e.g. 0.7% error rate in amplicons sequenced using R10.4.1 flowcell compared with 0.1% for Illumina) (10). While multiple tools exist for analysis of common amplicon data such as bacterial 16S rRNA sequencing (11–14), there is limited availability of tools for classifying arbitrary amplicons especially at resolution below bacterial species level, such as serovars of *Salmonella enterica*. AmpliconTyper is a flexible tool that can be used to identify ONT reads from a specific target organism in environmental or other samples with multiple diverse organisms, and to detect epidemiologically and phenotypically important alleles.

### Theory and implementation

#### Overview

AmpliconTyper is designed to differentiate ONT amplicon sequencing reads originating from amplification of target sequences (target organism) from off-target amplification (non-target organism). Non-target organisms are organisms other than the target organism that, in the user’s determination, may be amplified by the PCR primers used. The AmpliconTyper has two main functions: train and classify (**Fig. 1**). The former trains a machine learning (ML) model to distinguish ONT sequencing reads from target vs non-target organisms (**Fig. 2**). The latter classifies every read in an input file (FASTQ or BAM format) (15, 16) as target or non-target organism based on this model, and generates a report summarising the results (**Fig. 3**) (15, 16). Users can also supply AmpliconTyper with a variant call format (VCF) file (17) containing a list of genotypes and AMR-associated SNPs which, if identified in the data, are listed in the report (17). Additionally, if amplicons targeting accessory genes are included in the sequencing assay (e.g. acquired AMR genes), AmpliconTyper can call these as present/absent or with allelic variation, to support subtyping of the detected target pathogen. The version described here is v0.1.28 (DOI: 10.5281/zenodo.14621928).

**Figure 1.**
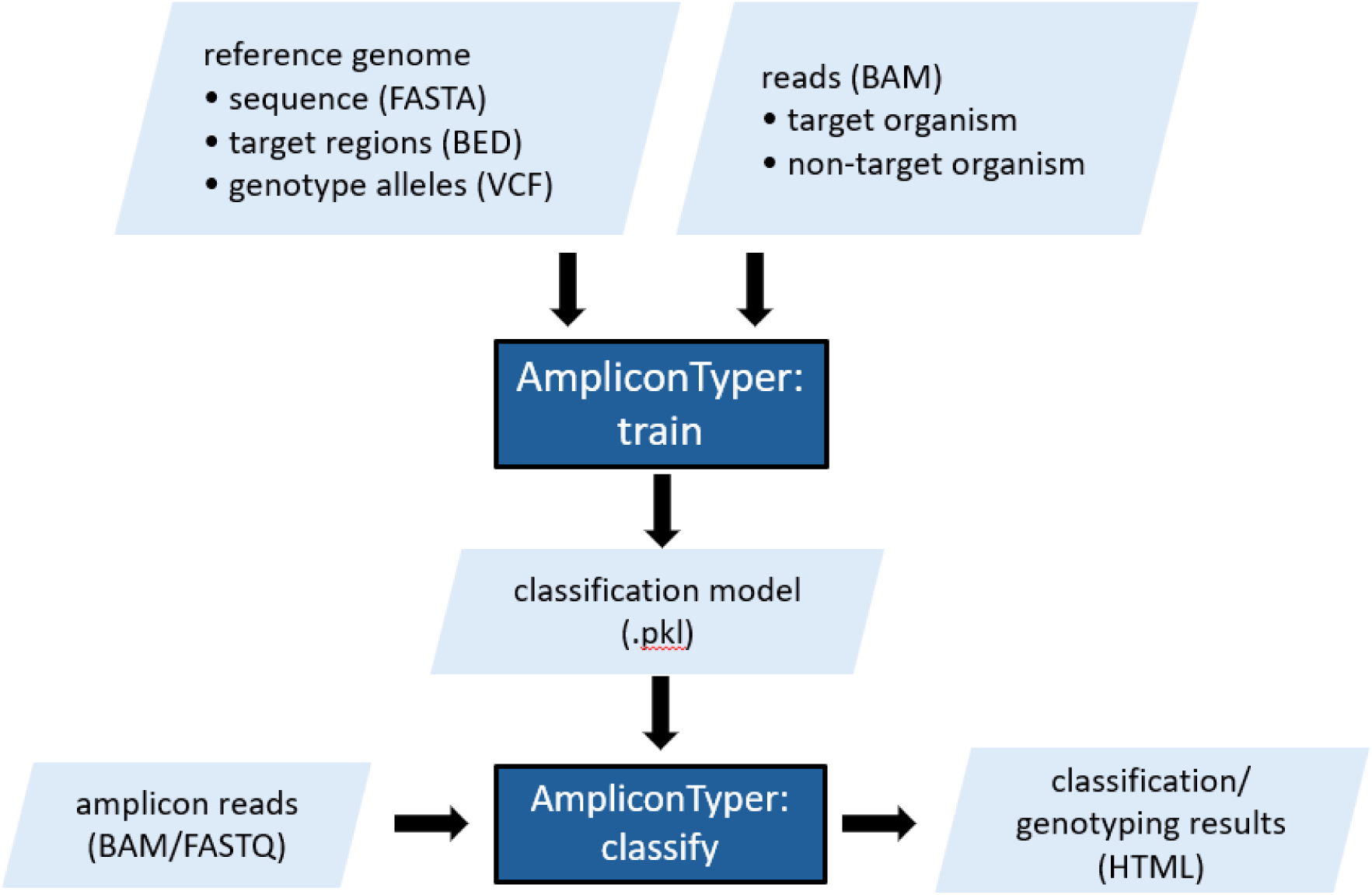
General workflow of the training and classification.

**Figure 2.**
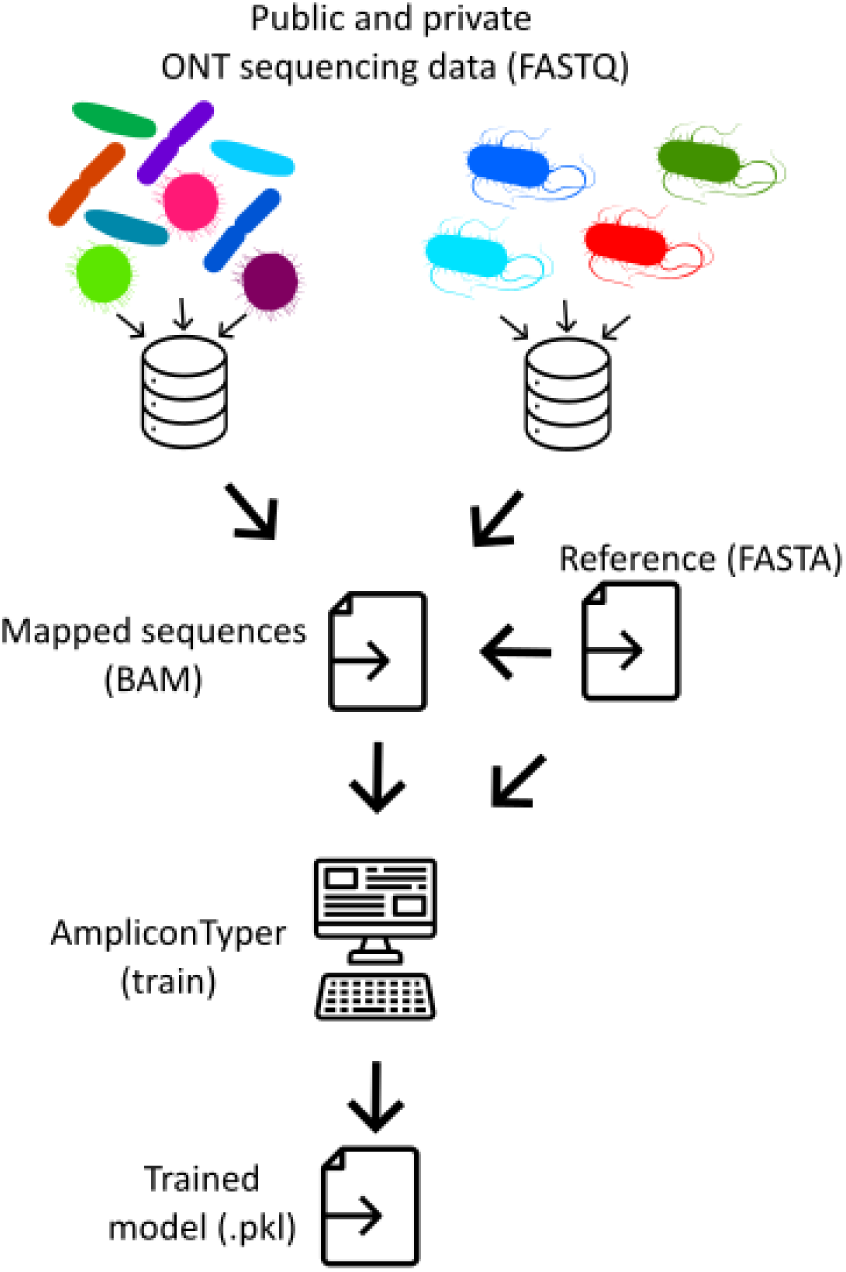
Overview of the training process. FASTQ files of WGS data from public and/or user databases are aligned to the desired amplicon sequences. The resulting BAM files, representing target and non-target organism sequences, are fed to the AmpliconTyper “train” function, which trains a machine learningclassifier and outputs it to a model file (.pkl) that can be used to classify new data (see Figure 2).

**Figure 3.**
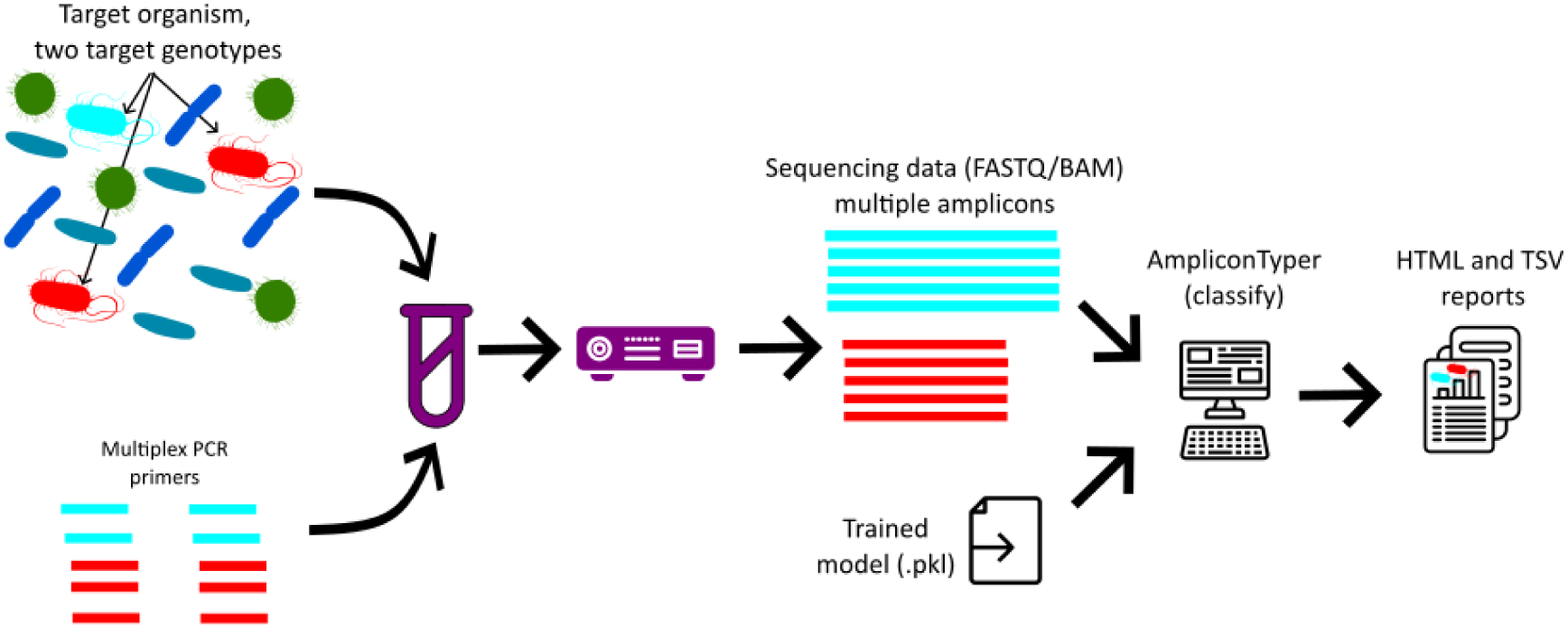
Genotyping workflow. The sample DNA is amplified in a single multiplex PCR. The resulting amplicons are sequenced on an ONT device and the sequence reads are analysed using AmpliconTyper. The output is a report (**File 1**) describing amplification results for each primer, and summary of mutations in the PCR products.

#### Training

AmpliconTyper relies on a trained ML model to classify ONT amplicon sequencing data from mixed sources such as environmental samples. The training process is summarised in **Figure 2** and **Supplementary Figure 1**. To train the model, user needs to provide the corresponding function “*train*” with:

a) the reference sequences as FASTA file,
b) BED file specifying which regions in the reference sequence are expected to be amplified (18),
c) directory with alignment of ONT reads from non-target organisms in BAM format (15),
d) directory with alignment of ONT reads from pure genomic material of target organism in BAM format (these can be from single or multiple genotypes),
e) VCF file (17) with list of positions in reference sequences that should be excluded from training as well as any genotype-or AMR-associated positions(17),

Since AmpliconTyper is designed for applications such as environmental surveillance where samples contain a mixture of organisms, training data should reflect the likely diversity of organisms in the sample sources. For this reason, we recommend that users supply AmpliconTyper not only user-generated amplicon data, but also publicly available whole genome sequencing (WGS) ONT read data mapped to target sequences (**Fig. 2**). As WGS data will contain reads that only partially overlap the target amplicon (**Supp. Fig. 2**), it is essential to discard such reads from training otherwise the model simply learns that reads with large number of missing nucleotides are from non-target organisms. When applying the trained model to new data, we also discard reads with one or more missing nucleotides either at the start or end of the reference sequence; though this can be overridden. As a consequence of using only reads that fully span amplicons, the public WGS data will typically have many fewer usable reads than the mean depth of coverage (**Supp. Fig. 2**). For this reason, we recommend using as many public sequencing libraries as possible. For our exemplar work on *S*. Typhi we used 10,301 non-Typhi and 68 *S*. Typhi WGS ONT libraries available via NCBI’s SRA database supplemented with one *S*. Typhi and two *S*. Paratyphi libraries we generated (**Supp. Data 1**) (1).

The training uses a logistic regression model with stochastic gradient descent, implemented as SGDClassifier with log-loss function in scikit-learn v1.5.2 (19). To train the model, the classifier first converts BAM files into a binary matrix with five columns per each nucleotide in the amplicon reference sequence to accommodate all possible nucleotide calls (A, C, T, G and N). We chose to discard reads with ambiguous nucleotide codes (i.e., N) due to the complexity they introduce (20). This classifier achieved much better performance compared to other tested classifiers including ensemble approaches (**Fig. 4**). The outlier amplicon in **Fig. 4B** is the off-target amplification of a single gene in *S*. Paratyphi C, which was poorly represented in the training set. The training process consists of splitting the data into a training set (80% or 5,000 of reads whichever is lower) and test set (20% or 5,000 of reads whichever is lower) with sensitivity and specificity as the model evaluation parameters. Importantly, all positions specified in the input VCF file (**Supp. Fig. 1**) are ignored during both training and classification(17). This is because some positions in the amplicons include multiple alleles that define lineages or AMR linked mutations and their inclusion reduces reliability of the classification.

**Figure 4.**
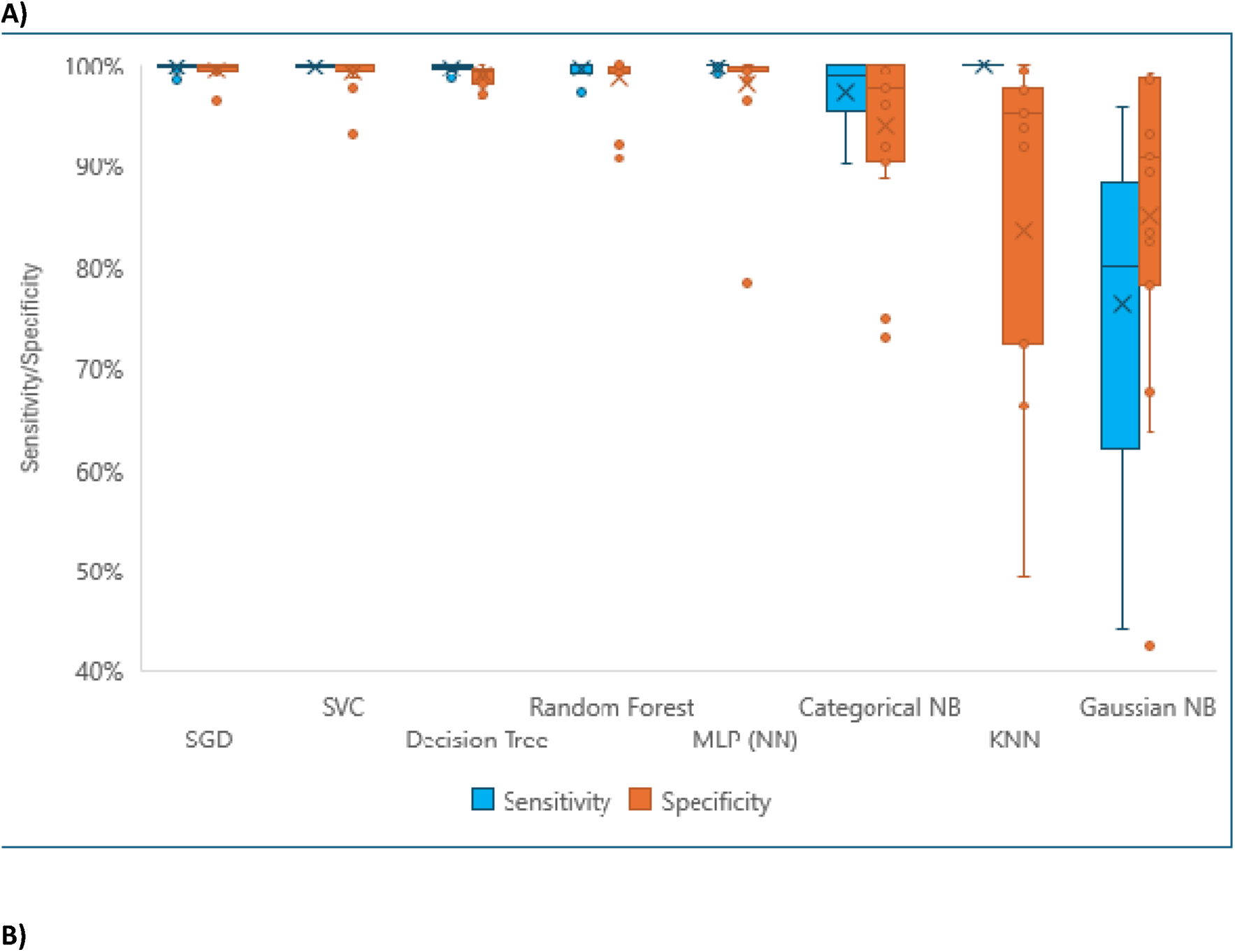
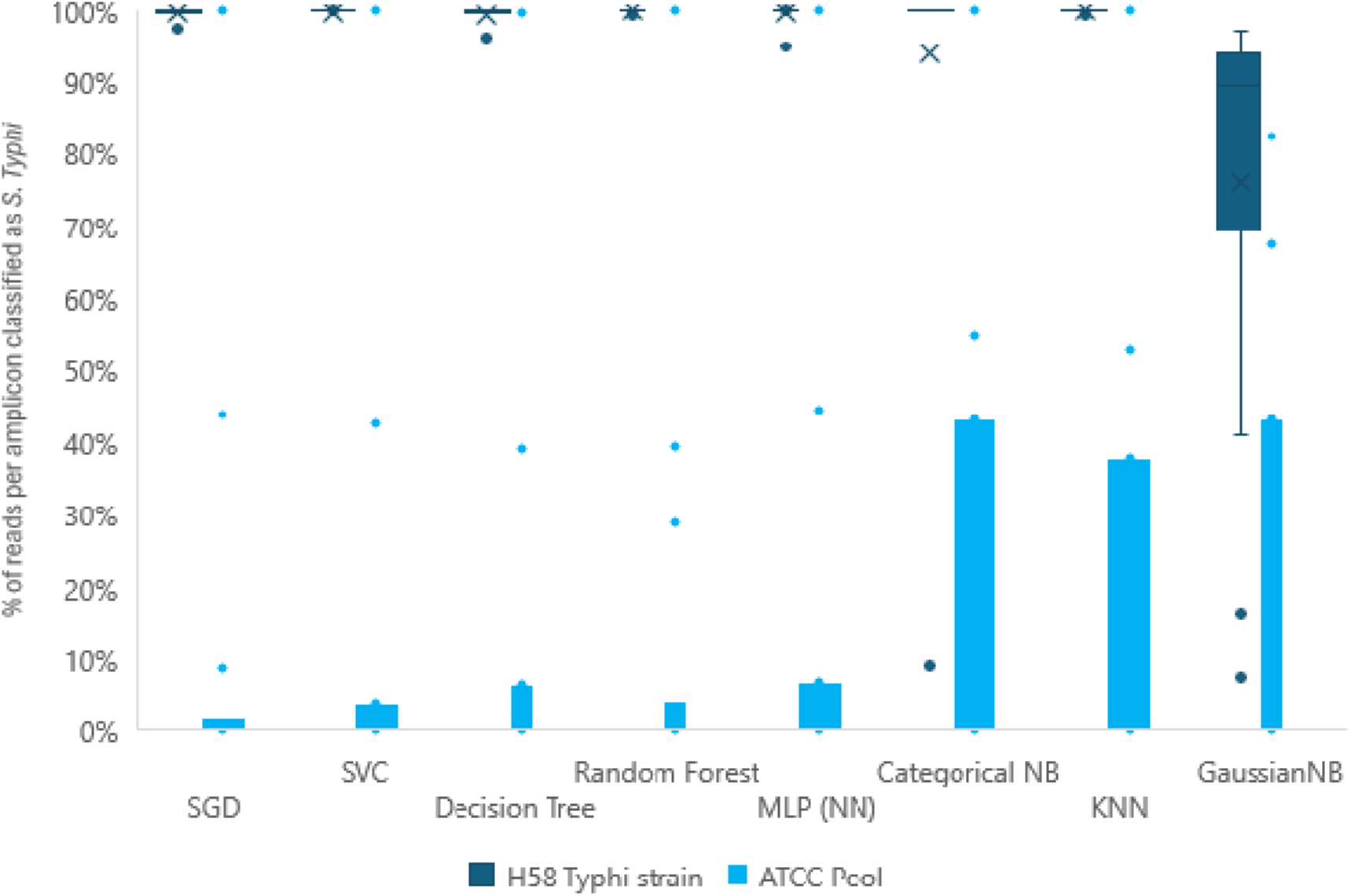
comparison of classification models. (A) The training dataset (mix of S. Typhi and non-S. Typhi ONT reads) was split into training data (80%) and test data (20%). Each classifier was trained on training data and evaluated against test data based on specificity and sensitivity. (B) Each trained classifier from (A) was tested against newly generated data consisting of one sample of S. Typhi strain H58 and one sample of pooled S. Paratyphi A, B and C, Aeromonas hydrophila, Klebsiella pneumoniae, and Citrobacter freundii from ATCC.

The above approach cannot deal with instances where the target organism sequence has no full-length homologs in non-target organisms, for example due to recombination events. For such cases, AmpliconTyper uses GaussianMixture model from scikit-learn package to determine the distribution of distances between the training target organism reads and the reference sequence (excluding known variable sites from the VCF). AmpliconTyper uses this distribution to classify new reads based on whether their distance from reference is within 10% of closest reads used in training. This approach delivers low specificity and sensitivity which are not *a priori* quantifiable. To mitigate this, when classifying new data, AmpliconTyper reports the nucleotide differences between reads classified as the target organism and the reference sequence.

Once the model is trained, it is saved into a Python object serialisation file (pickle file) that serves as input for the “*classify*” function. The pickle file (21) is a binary file that users cannot explore or modify other than through functions *“classify”*, *“train”* and *“genotyper_utilities”* provided with AmpliconTyper.

#### Classification of new data

The “*classify*” function applies the trained model to new data (**Fig. 3, Supp. Fig. 3**). It requires:

a) Model pickle file (generated during model training as described above),
b) Either FASTQ or BAM file with reads to classify (15, 16).

The classifier first maps the sequencing data to the reference sequence using minimap2 v2.1 (22) and then applies pre-trained models described above to each read in order to assign reads to one of the two categories: target organism or non-target organism. For each amplicon, the reads classified as target organism are used to generate a consensus sequence which is compared to the reference sequence and the differences are reported to the user. (Haplotyping functionality, to resolvemixtures of target-organism genotypes, is under development and will be included in a future version.)

We have included an option to assign genotype or AMR labels to specific alleles to facilitate interpretation of AmpliconTyper outputs for amplicon-based genotyping. The allele information is supplied during model training and is stored within the model. We have opted to use the ‘ID’ field of the VCF file to store this information (**Supp. Data 2**) (17). The value of the ID field is assumed to consist of two parts: allele association and allele type. For example, “4:GT” means alternative (ALT) allele at this site implies genotype (GT) 4, whereas “gyrA_D87G:AMR” means ALT allele implies AMR genotype gyrA_D87G. Positions with neither “:GT” nor “:AMR” suffix are treated as not informative in the AmpliconTyper genotype report.

The results of classification are written to an HTML formatted report (**File 1**) which provides the summary of results, the genotypes supported by the data, detailed per-sample report of mapping results, and identified differences from reference sequence/s.

While training function requires large volume of data, so providing test data is impractical, the classification functionality can be tested using provided test data (doi: 10.5281/zenodo.14621928). Once AmpliconTyper has been installed and test data extracted from archive, the classification function can be run using *classify -f ./test_data/fastqs/ -b ./test_data/bams/ -d./test_data/metadata.tsv -c Description -o test_report.html -m ./test_data/typhi_v8.pkl*.

### Performance

We used AmpliconTyper on a Linux server (64 GB RAM, 8 Intel Xeon E5-2640 CPUs), but model training can in principle be done on a mid-level desktop, and classification is designed to work on a mid-level desktop computer (**Table 1**). To benchmark the AmpliconTyper training function we used a reduced *S*. Typhi dataset which consists of n=2,000 BAM files with negative training data, 6 GB of BAM files with positive data. To benchmark alternative algorithms for the classification step (**Fig. 4**) we used 8 samples of multiplex amplicon sequencing of *S*. Typhi genomic DNA with 904,795 mapped reads (241 MB).

**Table 1.** performance of training and classification on different computers.

|  | Server (8 CPUs) | Desktop (1 CPU) |
| --- | --- | --- |
| CPU type | Intel Xeon E5-2640 | Intel i3-7100 |
| Operating system | Linux | Linux ran via Windows WSL |
| Training data loading* | ~10 min | ~1.2 hr |
| Model training* | ~2 min | ~2 min |
| Memory peak | ~2.5 GB | ~0.5 GB |
| Classification of 230 MB of new data* | ~2.5 min | ~2 min |

\* - elements which are parallelised.

## Discussion

AmpliconTyper offers a simple way to detect epidemiologically relevant markers for a target organism from ONT amplicon sequencing of environmental or similarly contaminated samples. The AmpliconTyper is organism agnostic, relies on standard input file formats and can enrich output with identification of specific genotype and AMR alleles. A separate tool, AmpliSeqDesigner (https://github.com/AntonS-bio/AmpliSeqDesigner), was also developed to design the primers used in this study. However, the two pipelines work independently, and AmpliconTyper can work with primers designed using other tools.

AmpliconTyper uses standard input files such as FASTA, VCF, BED and BAM that should be familiar to potential users (15, 17, 18). The output, while simple in appearance, is informative and gives the user both a quick summary and detailed results.

For installation of the AmpliconTyper we strongly encourage using a package manager (conda or mamba) to install the tool from Bioconda, but AmpliconTyper can also be installed directly from its GitHub repository (https://github.com/AntonS-bio/AmpliconTyper). Once installed, the tool exposes two commands: “*train*” and “*classify*” both of which are meant to be run via command line. The tool can be used with either Linux or MacOS, and Windows 10 or higher (via the Windows Subsystem Linux).

The main limitation of the AmpliconTyper is that it classifies the reads into two categories: target and non-target. While we plan to introduce support for multiple classification classes in the future, currently multiple organisms must be modelled separately.

Another limitation stems from the requirement for reliable training data, with sufficient high-quality data confirmed as target and non-target. When public data is relied on, care must be taken to assess the quality of the sequence data, and of the target/non-target labels. To assist users to identify issues with the taxonomic labelling of training data, the training AmpliconTyper function outputs a list of samples with misclassified reads and the number of reads from each that were misclassified.

## Conclusions

AmpliconTyper offers a dedicated tool for analysis of multiplex ONT amplicon sequencing data from environmental or other samples with multiple organisms present. It relies on standard genomic data formats, provides a comprehensive results report, supports inference of genotype and AMR from alleles in the data, and permits model training using public or user-generated data. AmpliconTyper fills an important gap in the bioinformatics toolkit, and we look forward to continuing its improvement with community feedback.

## Abbreviations

AMR: antimicrobial resistance
ATCC: American type culture collection
BAM: binary alignment map
CPU: central processing unit
GB: gigabytes
HTML: hypertext markup language
KNN: k-nearest neighbors
MB: megabytes
ML: Machine Learning
MLP: Multi-layer perceptron
NB: naïve Bayes
NCBI: National Center for Biotechnology Information
ONT: Oxford Nanopore Technologies
PCR: polymerase chain reaction
SGD: stochastic gradient descent
SVC: support vector classifier
VCF: Variant Call Format
WGS: whole genome sequencing

## Availability and requirements

Project name: AmpliconTyper

Project home page: https://github.com/AntonS-bio/AmpliconTyper.git

Operating system(s): Linux, MacOS or Windows 10+ with Windows Subsystem Linux

Programming language: Python v3.10.12

Other requirements: None

License: GNU GPL v3

## Ethics approval and consent to participate

Not applicable.

## Consent for publication

Not applicable.

## Authors’ contributions

K.H., N.G and Z.A.D. conceived the idea and designed the study. A.S. developed the software. J.M. and C.T. generated DNA sequencing data for analysis and tested the software. All authors read and approved the final manuscript.

---

**Supplementary Figure 1.**
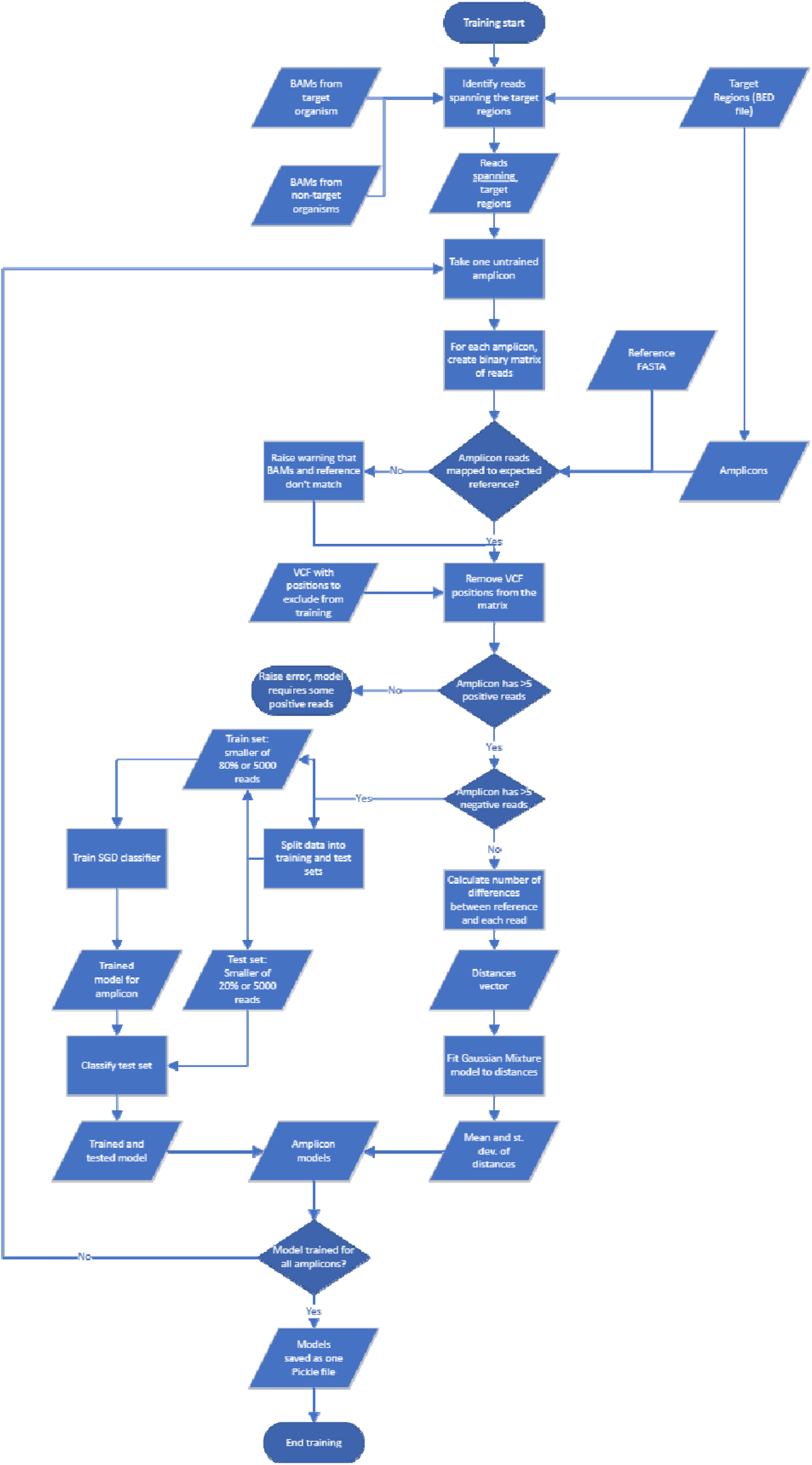
Training workflow.

**Supplementary Figure 2.**
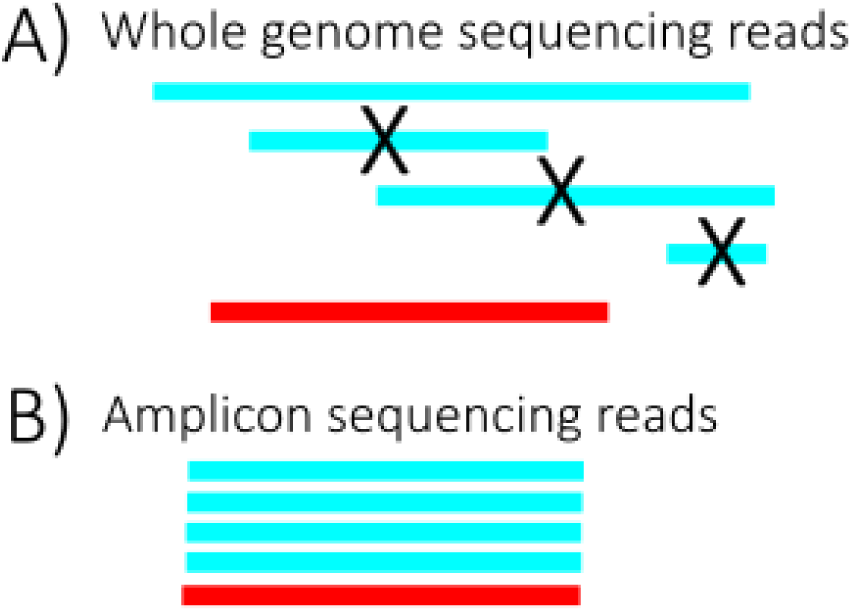
Stylised distribution of reads (blue) against target amplicon (red) in whole genome and amplicon sequencing. Only reads that fully span the target amplicon are used for model training.

**Supplementary Figure 3.**
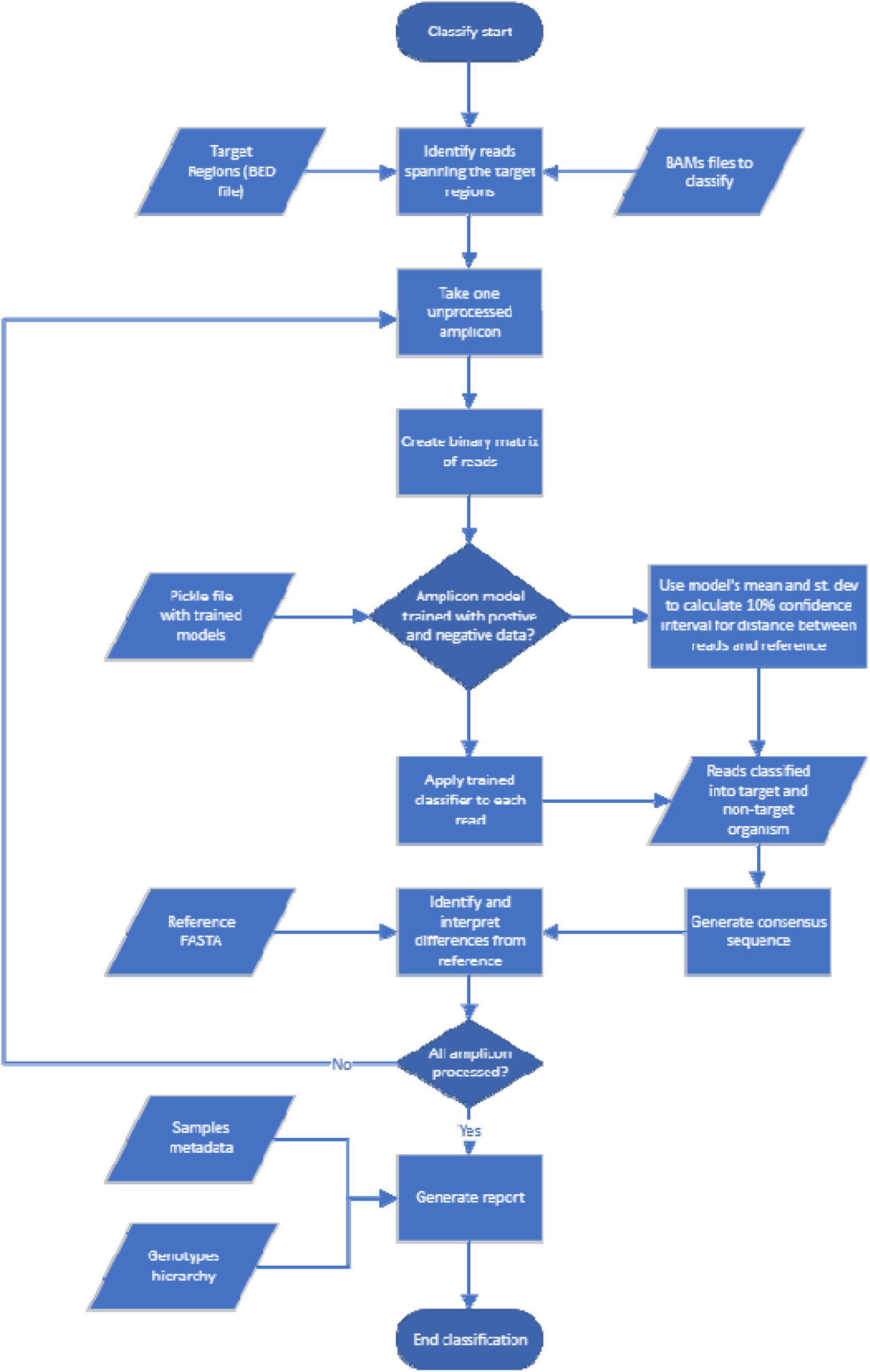
Classification work flow.

## Summary of results

Model name: typhi_v8, model date: 18/Dec/24

| Sample ? | Amplicons by # of dominant unknown SNPs ? | Possible High Level Genotypes ? | Implied Genotypes ? | Implied AMR ? | Sequencing Reads ? | Mapped, correct length ? | Mapped, but too short/long ? |
| --- | --- | --- | --- | --- | --- | --- | --- |
| barcode1 S. Typhi negative wastewater | 2, 0, 0, 0, 0, 0 | - | - | - | 32167 | 3015 | 820 |
| barcode2 S. Typhi H58 + wastewater | 11, 1, 0, 0, 0, 0 | 4 | 4.3.1.1 (94%) | gyrA_S83T (99%) | 14187 | 2534 | 983 |

"-" fewer than 2 amplicons with 50 target organism reads and fewer than 2 SNPs
Classifier allowed soft clip (-s) of 25nt and 10nt length difference (-l) between reference and mapped read.
Report generated on 20/Dec/24 at 12:38. See models list

### barcode 1

**S. Typhi negative wastewater**

**Back to Summary**

| Amplicon ? | Tot. Reads ? | Rejected on Length ? | % Target Org. ? | # Target Org. ? | All known SNPs ? | Dominant known SNPs ? | Dominant unknown SNPs ? | Implied Genotypes ? | Implied AMR ? |
| --- | --- | --- | --- | --- | --- | --- | --- | --- | --- |
| 1_3_4.3.1 | * | - | - | - | - | - | - | - | - |
| 2.2.2_2 | * | - | - | - | - | - | - | - | - |
| 2.3.1_v12 | * | - | - | - | - | - | - | - | - |
| 2.3.2 | 2215 | 447 | 99.66% | 1762 | 0 | 0 | 0 |  |  |
| 2.5_4.3.1.2.1_3.3 | * | - | - | - | - | - | - | - | - |
| 3.1.1_v5 | * | - | - | - | - | - | - | - | - |
| 3.3.1 | * | - | - | - | - | - | - | - | - |
| 4.3.1.1 | * | - | - | - | - | - | - | - | - |
| 4.3.1.1.P1_v2 | * | - | - | - | - | - | - | - | - |
| 4.3.1.2.1.1_4 | * | - | - | - | - | - | - | - | - |
| 4.3.1.2_v3 | * | - | - | - | - | - | - | - | - |
| parC_v7 | * | - | - | - | - | - | - | - | - |
| tviD_v2 | * | - | - | - | - | - | - | - | - |
| acrB_v2 | 1463 | 311 | 99.83% | 1150 | 0 | 0 | 0 |  |  |
| gyrA | * | - | - | - | - | - | - | - | - |
| plasmid_ST6_C19241A^ | * | - | - | - | - | - | - | - | - |
| chromosomal_MDR_cyaA | * | - | - | - | - | - | - | - | - |
| chromosomal_MDR_yidA | * | - | - | - | - | - | - | - | - |
| plasmid_MDR_AL513383.1 | * | - | - | - | - | - | - | - | - |
| plasmid_MDR_CP029645.1 | * | - | - | - | - | - | - | - | - |
| plasmid_MDR_CP029879.1 | * | - | - | - | - | - | - | - | - |
| plasmid_MDR_CP029924.1 | * | - | - | - | - | - | - | - | - |
| plasmid_MDR_CP029957.1 | * | - | - | - | - | - | - | - | - |
| plasmid_MDR_LT904879.1 | * | - | - | - | - | - | - | - | - |
| plasmid_MDR_LT904892.1 | * | - | - | - | - | - | - | - | - |

\* Amplicon has fewer than 51 target organism reads

### barcode 2

**S. Typhi H58+ wastewater**

**Back to Summary**

| Amplicon ? | Tot. Reads ? | Rejected on Length ? | % Target Org. ? | # Target Org. ? | All known SNPs ? | Dominant known SNPs ? | Dominant unknown SNPs ? | Implied Genotypes ? | Implied AMR ? |
| --- | --- | --- | --- | --- | --- | --- | --- | --- | --- |
| 1_3_4.3.1 | 143 | 65 | 100.00% | 78 | 1 | 1 | 0 | 4.3.1 (96%) |  |
| 2.2.2_2 | 212 | 112 | 100.00% | 100 | 0 | 0 | 0 |  |  |
| 2.3.1_v12 | 191 | 50 | 100.00% | 141 | 1 | 0 | 0 | 2.3.1 (18%) (?) |  |
| 2.3.2 | 555 | 103 | 100.00% | 452 | 0 | 0 | 0 |  |  |
| 2.5_4.3.1.2.1_3.3 | 326 | 111 | 99.53% | 214 | 0 | 0 | 0 |  |  |
| 3.1.1_v5 | * | - | - | - | - | - | - | - | - |
| 3.3.1 | 150 | 37 | 99.12% | 112 | 0 | 0 | 0 |  |  |
| 4.3.1.1 | 201 | 39 | 100.00% | 162 | 1 | 1 | 0 | 4.3.1.1 (94%) |  |
| 4.3.1.1.P1_v2 | 72 | 13 | 100.00% | 59 | 0 | 0 | 0 |  |  |
| 4.3.1.2.1.1_4 | 433 | 186 | 99.19% | 245 | 1 | 1 | 0 | 4 (99%) |  |
| 4.3.1.2_v3 | * | - | - | - | - | - | - | - | - |
| parC_v7 | * | - | - | - | - | - | - | - | - |
| tviD_v2 | 392 | 92 | 99.33% | 298 | 1 | 1 | 0 | known, but uninformative allele (95%) |  |
| acrB_v2 | 605 | 93 | 99.61% | 510 | 0 | 0 | 0 |  |  |
| gyrA | 157 | 64 | 100.00% | 93 | 1 | 1 | 1 |  | gyrA_S83 T (99%) |
| plasmid_ST6_C19241A^ | * | - | - | - | - | - | - | - | - |
| chromosomal_MDR_cyaA | * | - | - | - | - | - | - | - | - |
| chromosomal_MDR_yidA | * | - | - | - | - | - | - | - | - |
| plasmid_MDR_AL513383.1 | * | - | - | - | - | - | - | - | - |
| plasmid_MDR_CP029645.1 | * | - | - | - | - | - | - | - | - |
| plasmid_MDR_CP029879.1 | * | - | - | - | - | - | - | - | - |
| plasmid_MDR_CP029924.1 | * | - | - | - | - | - | - | - | - |
| plasmid_MDR_CP029957.1 | * | - | - | - | - | - | - | - | - |
| plasmid_MDR_LT904879.1 | * | - | - | - | - | - | - | - | - |
| plasmid_MDR_LT904892.1 | * | - | - | - | - | - | - | - | - |

\* Amplicon has fewer than 51 target organism reads

### Consensus sequence

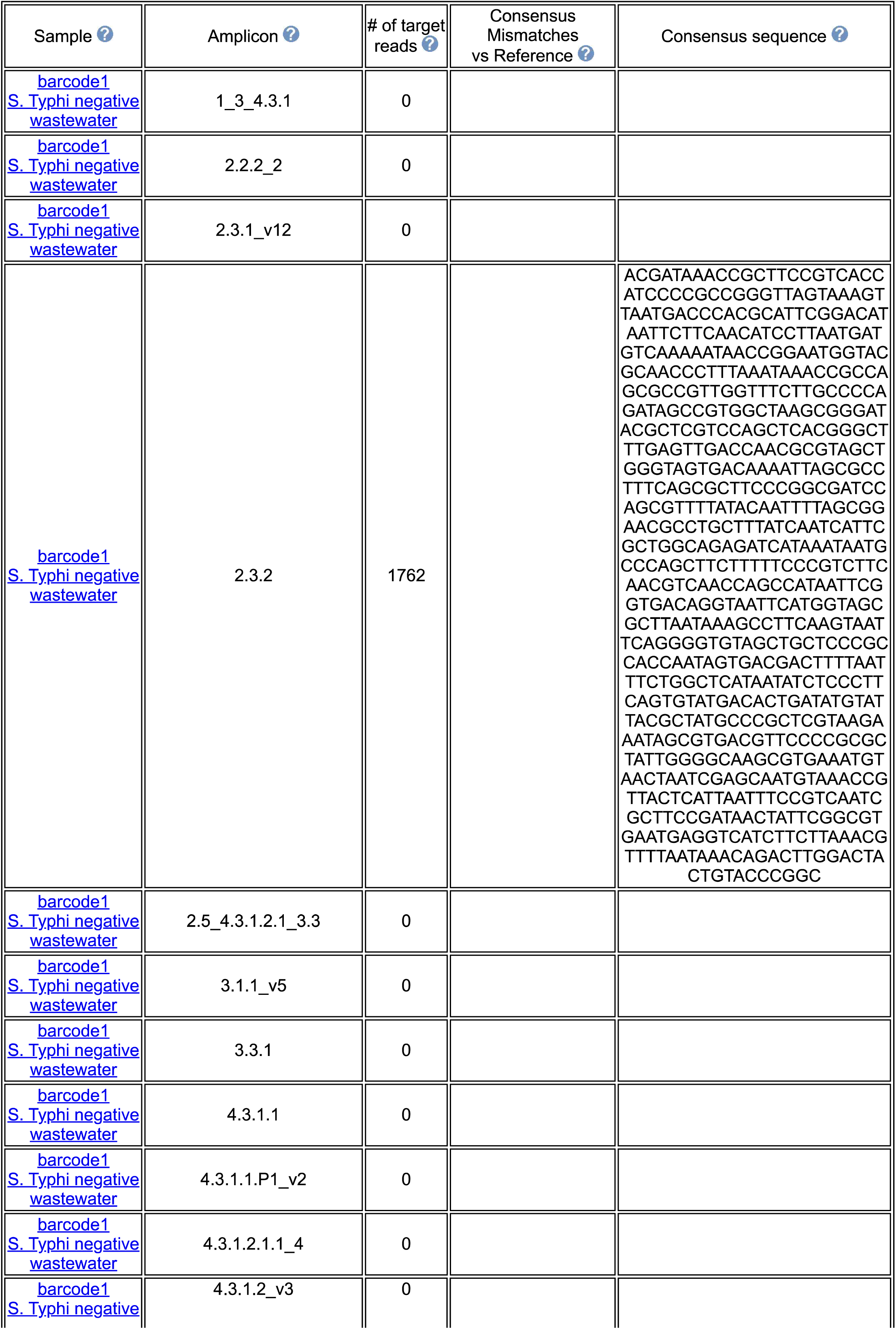
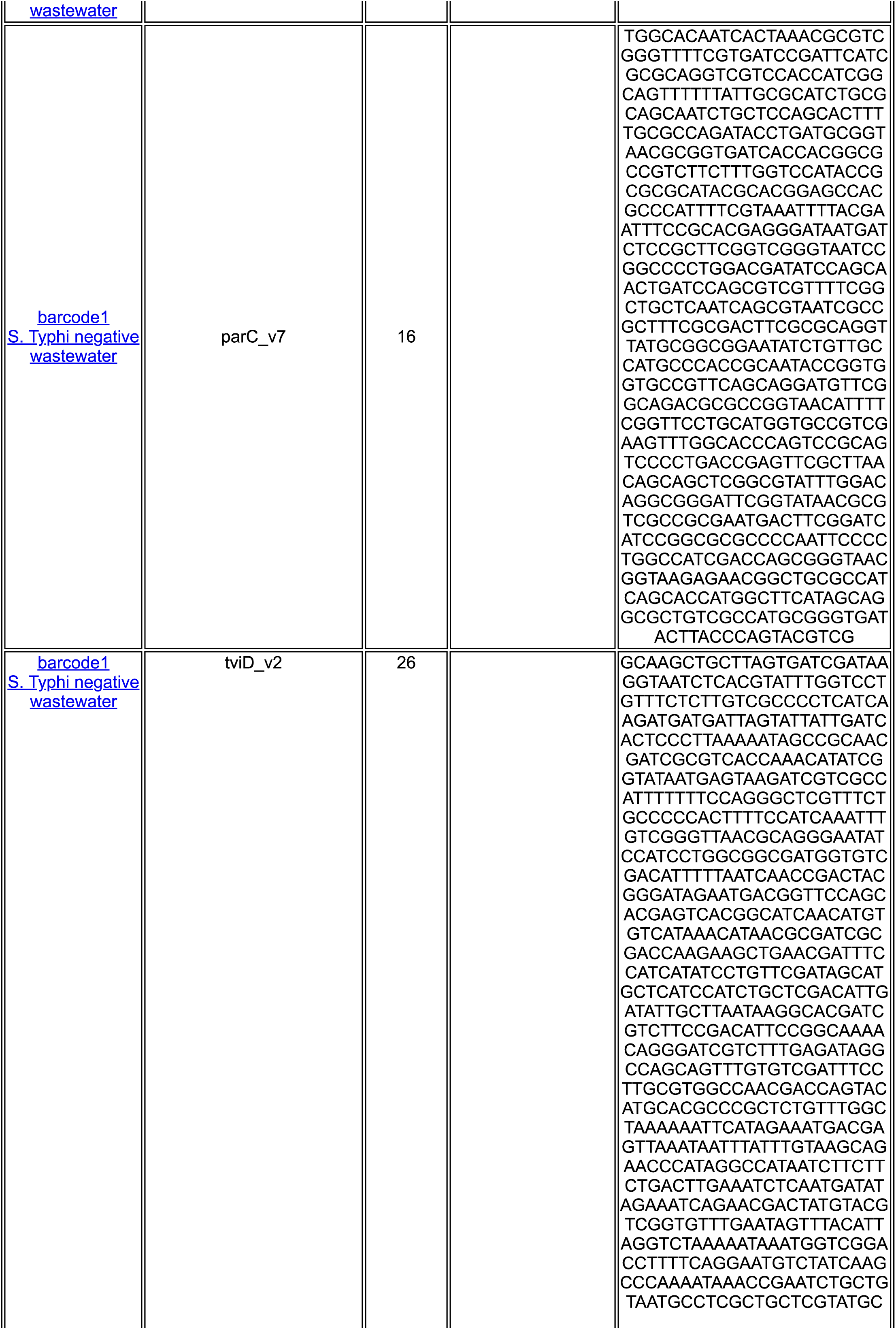
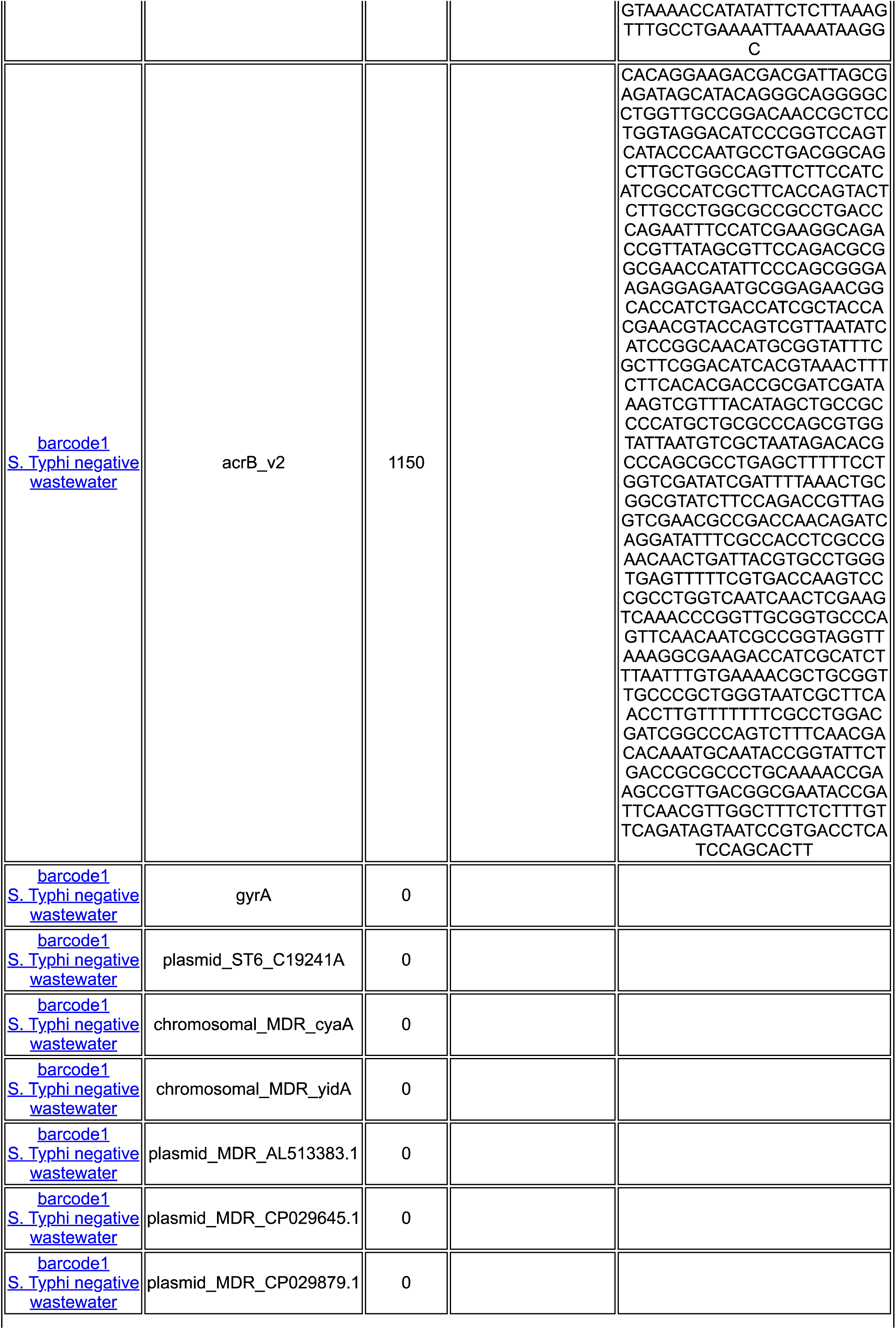
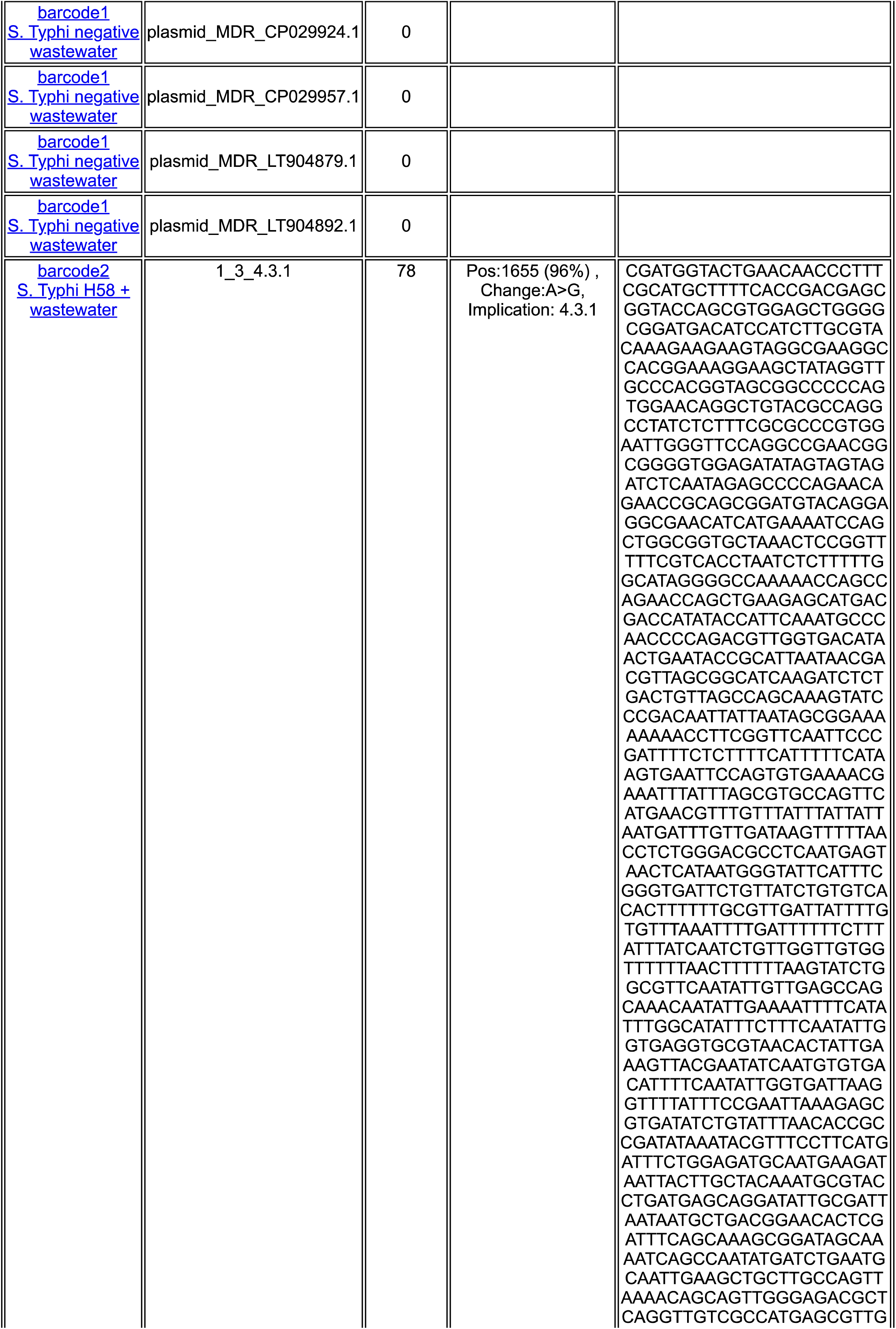
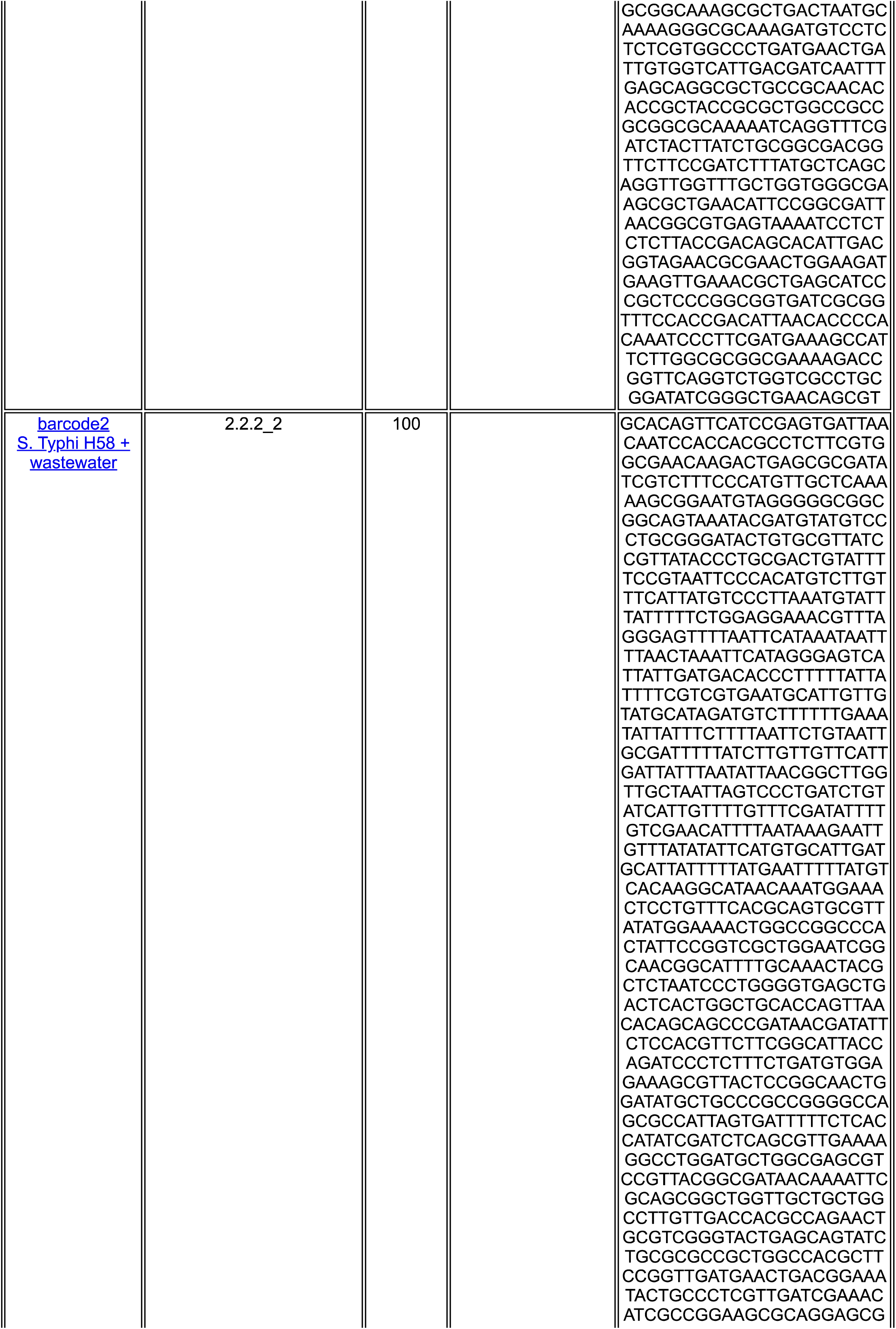
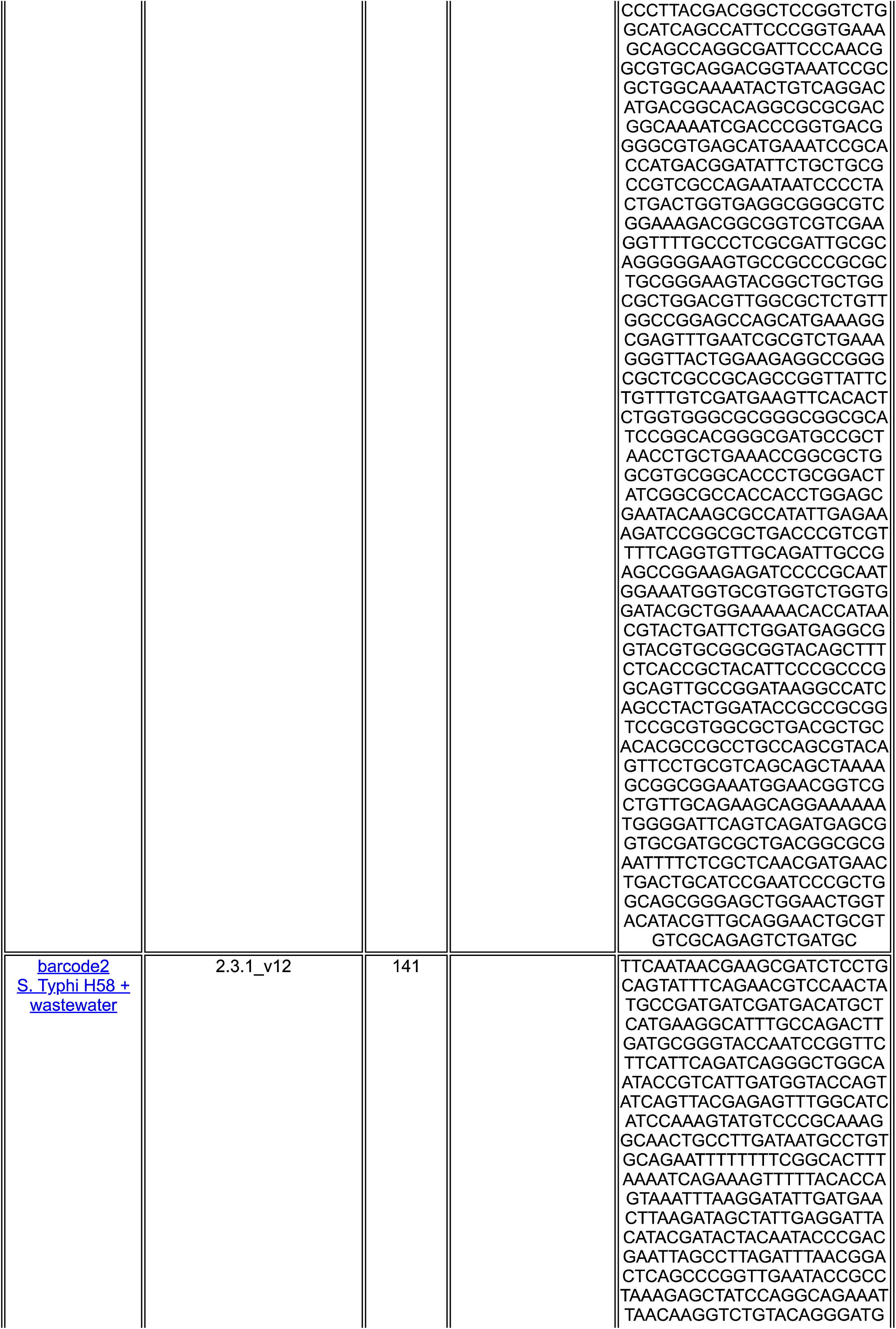
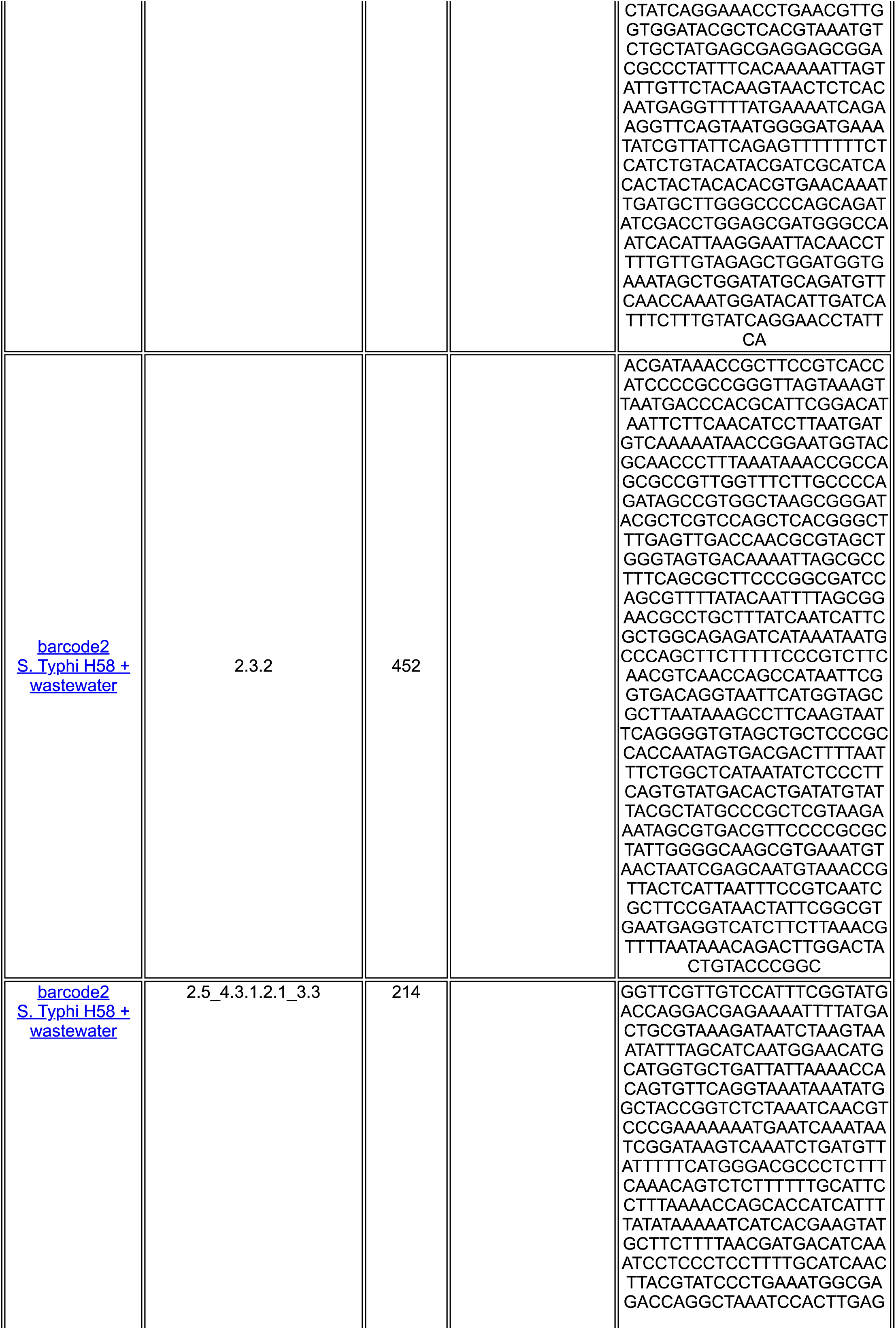
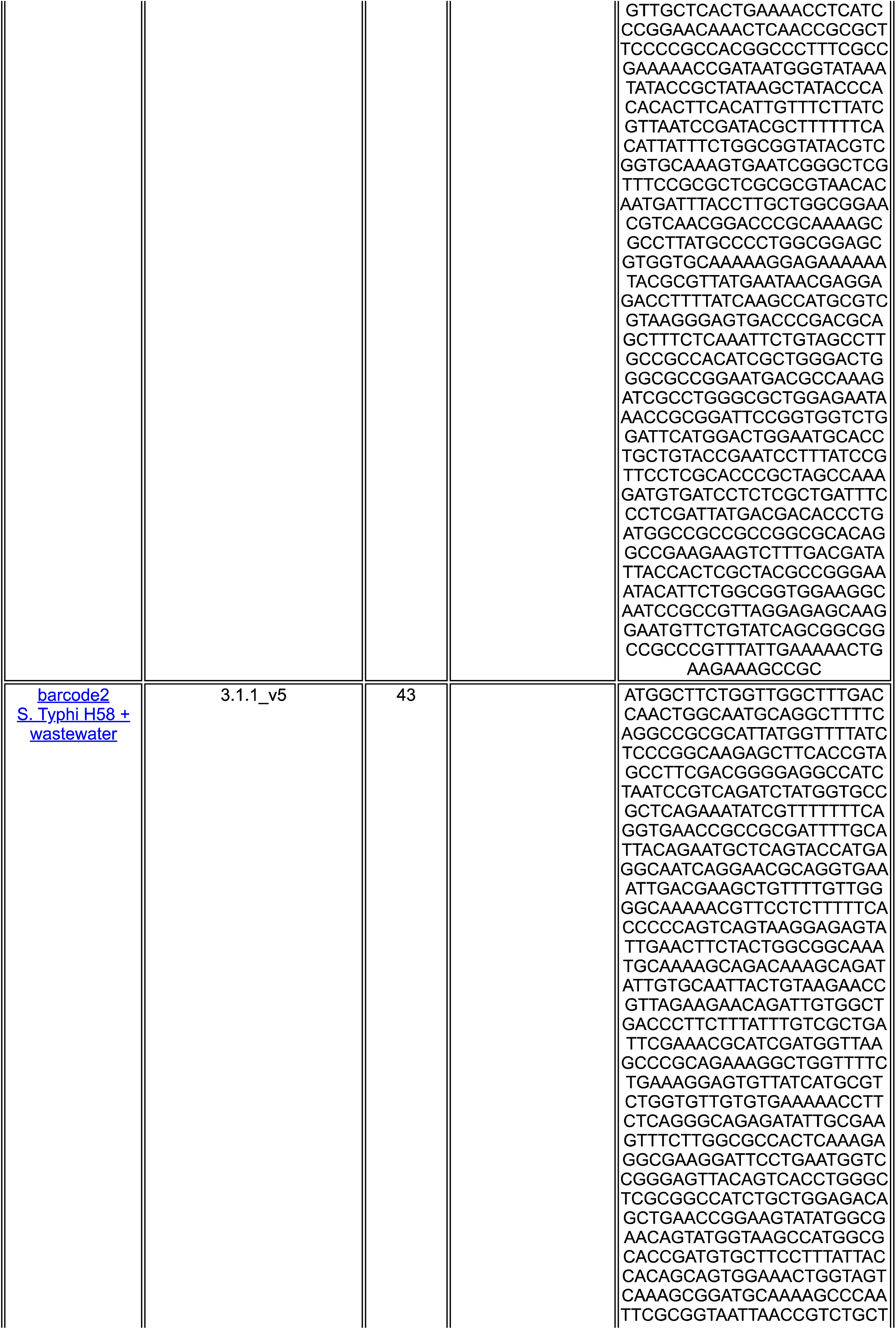
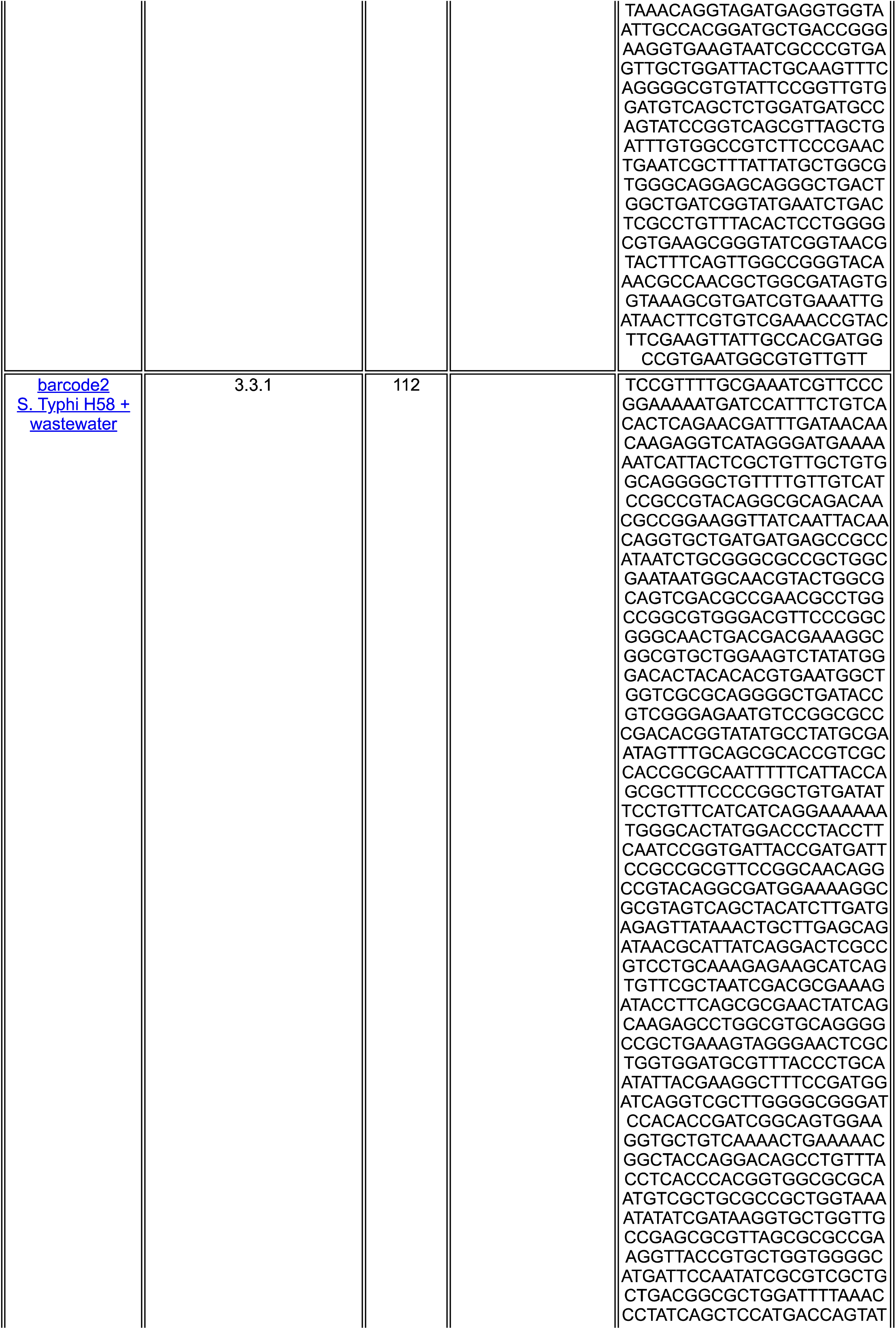
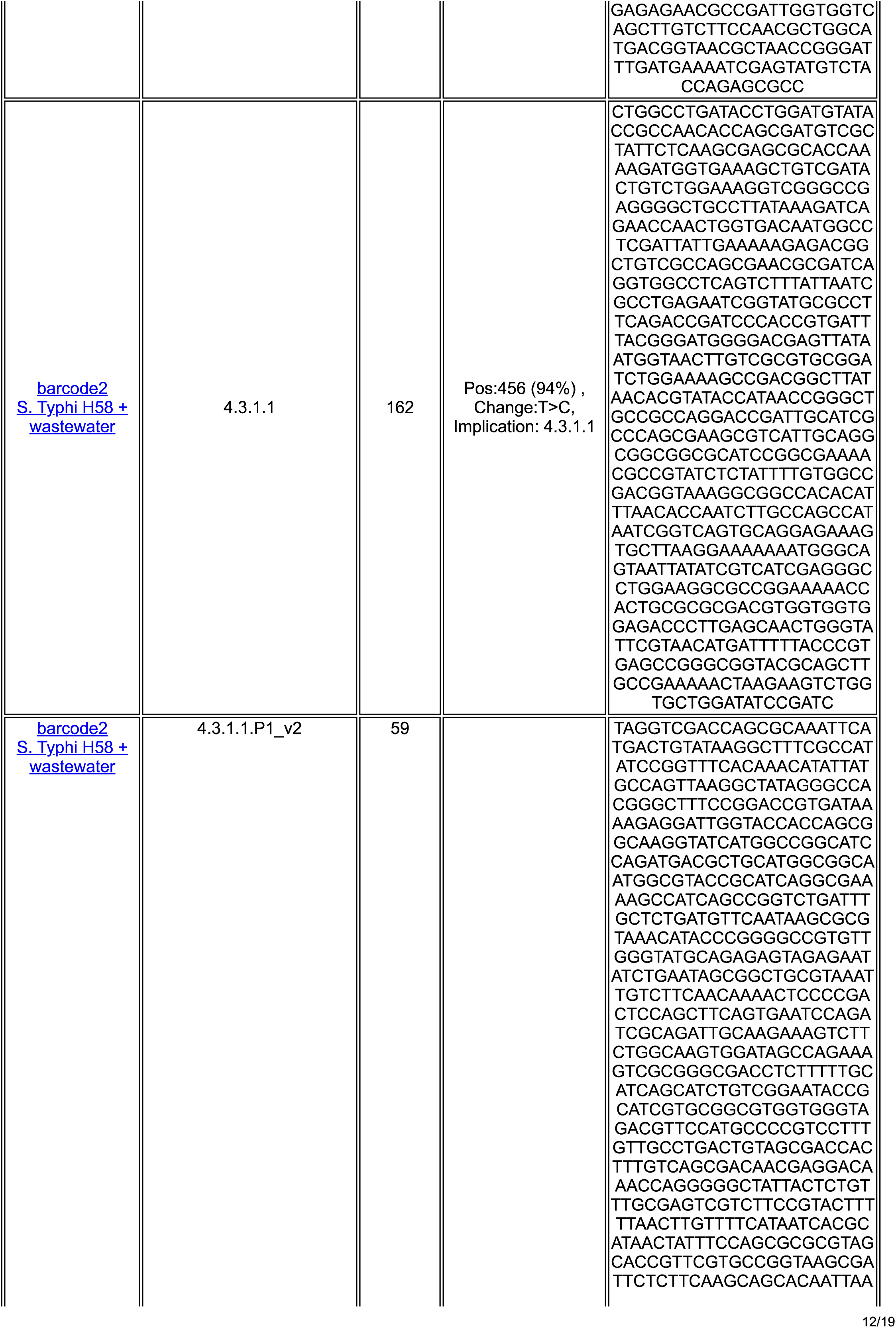
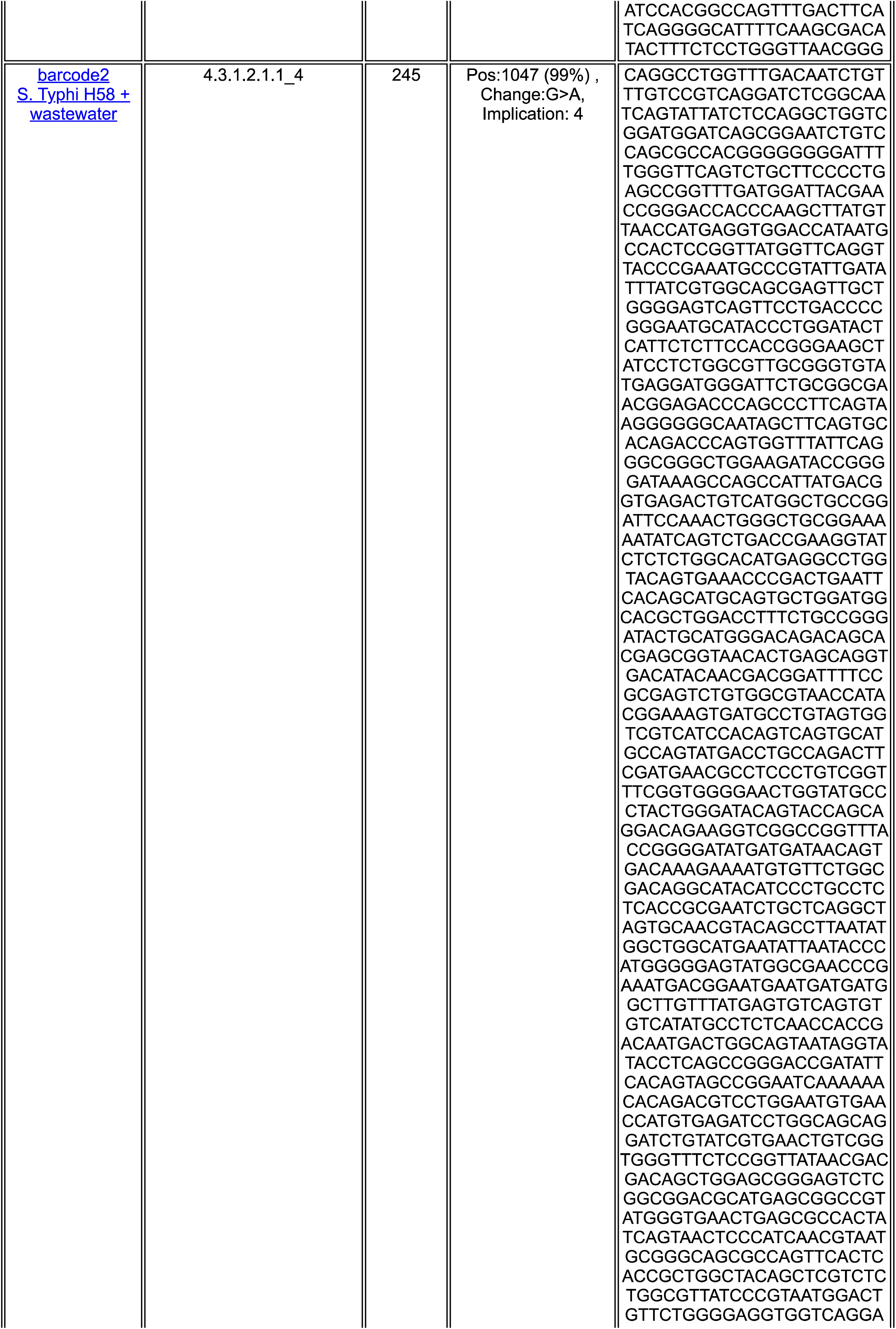
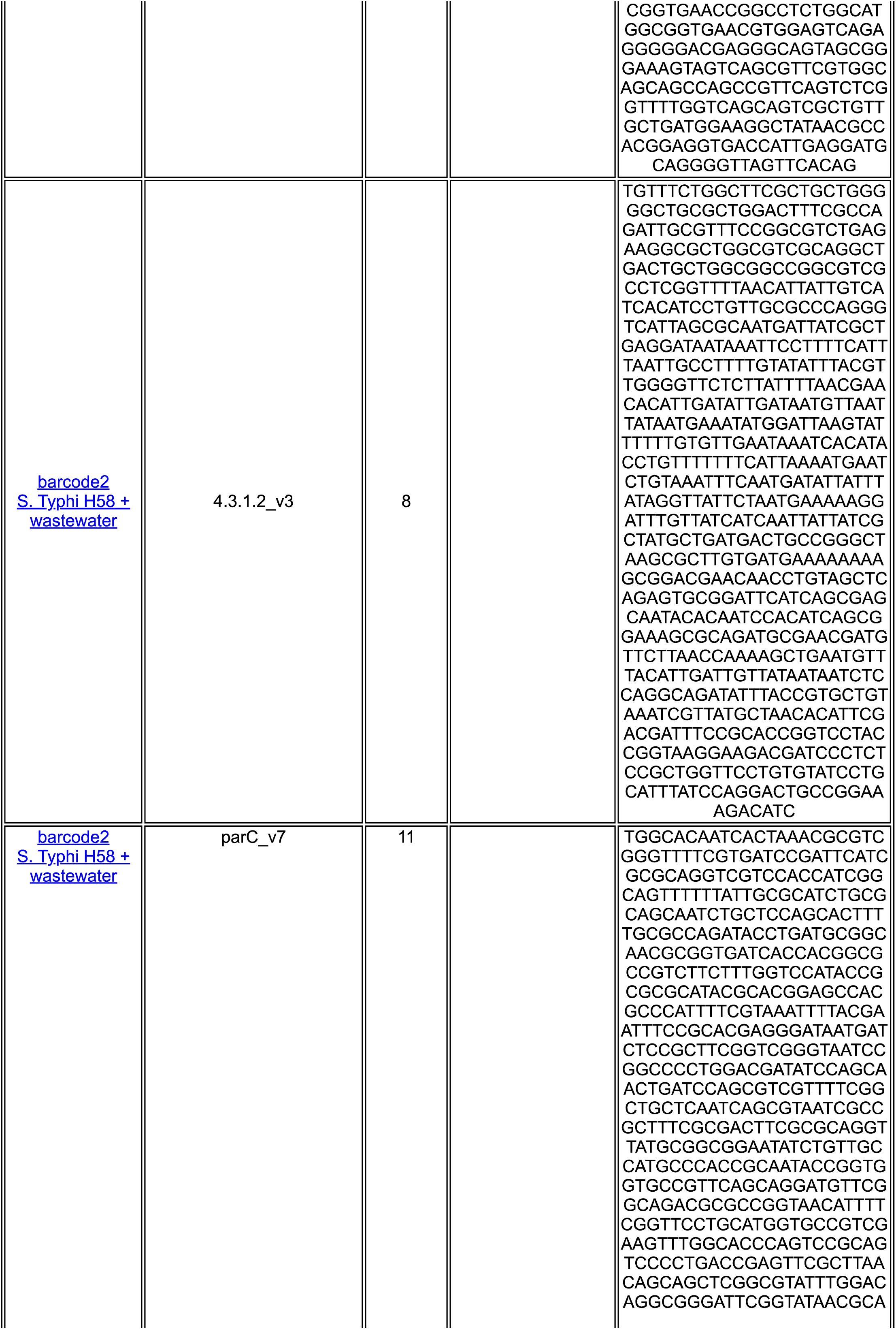
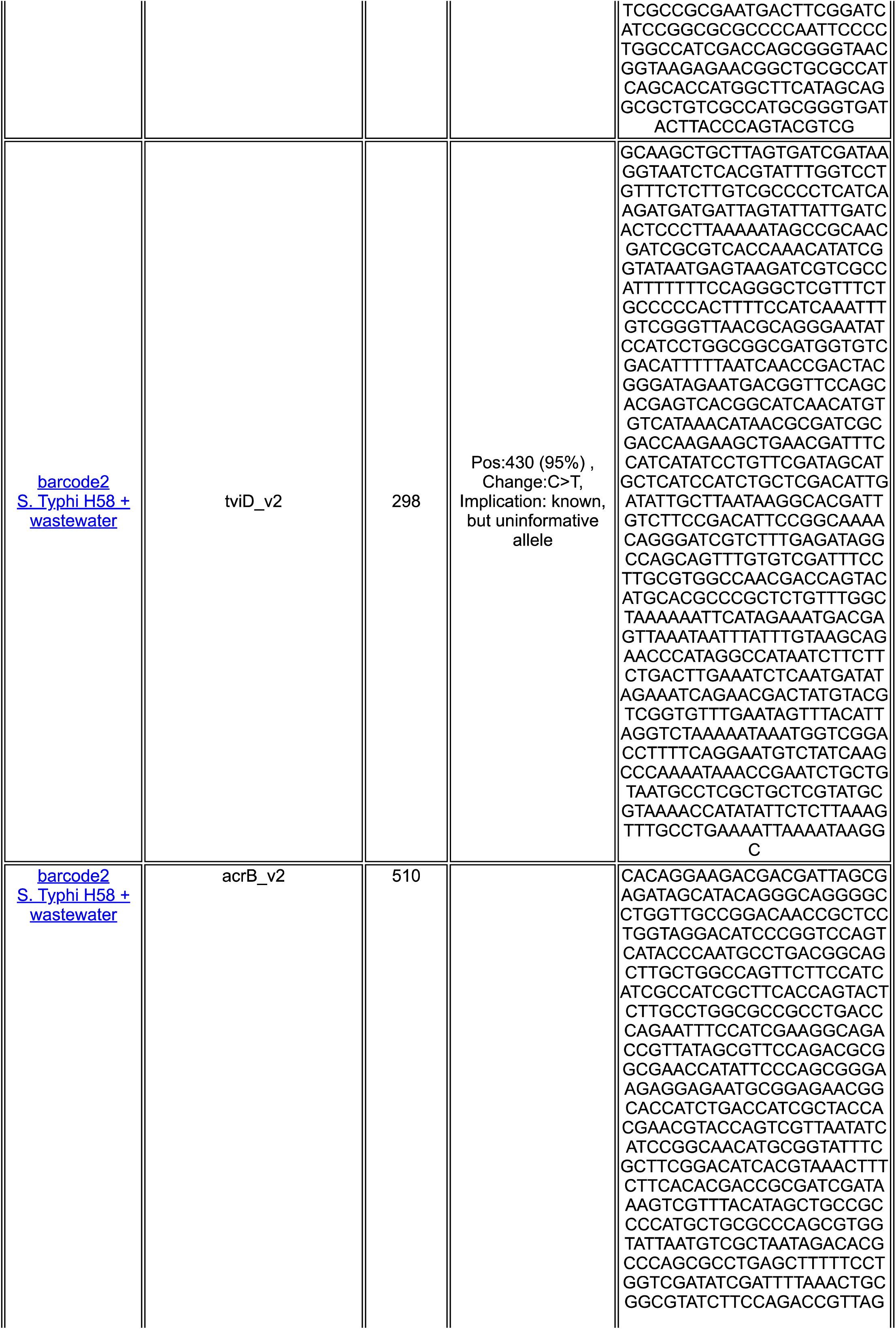
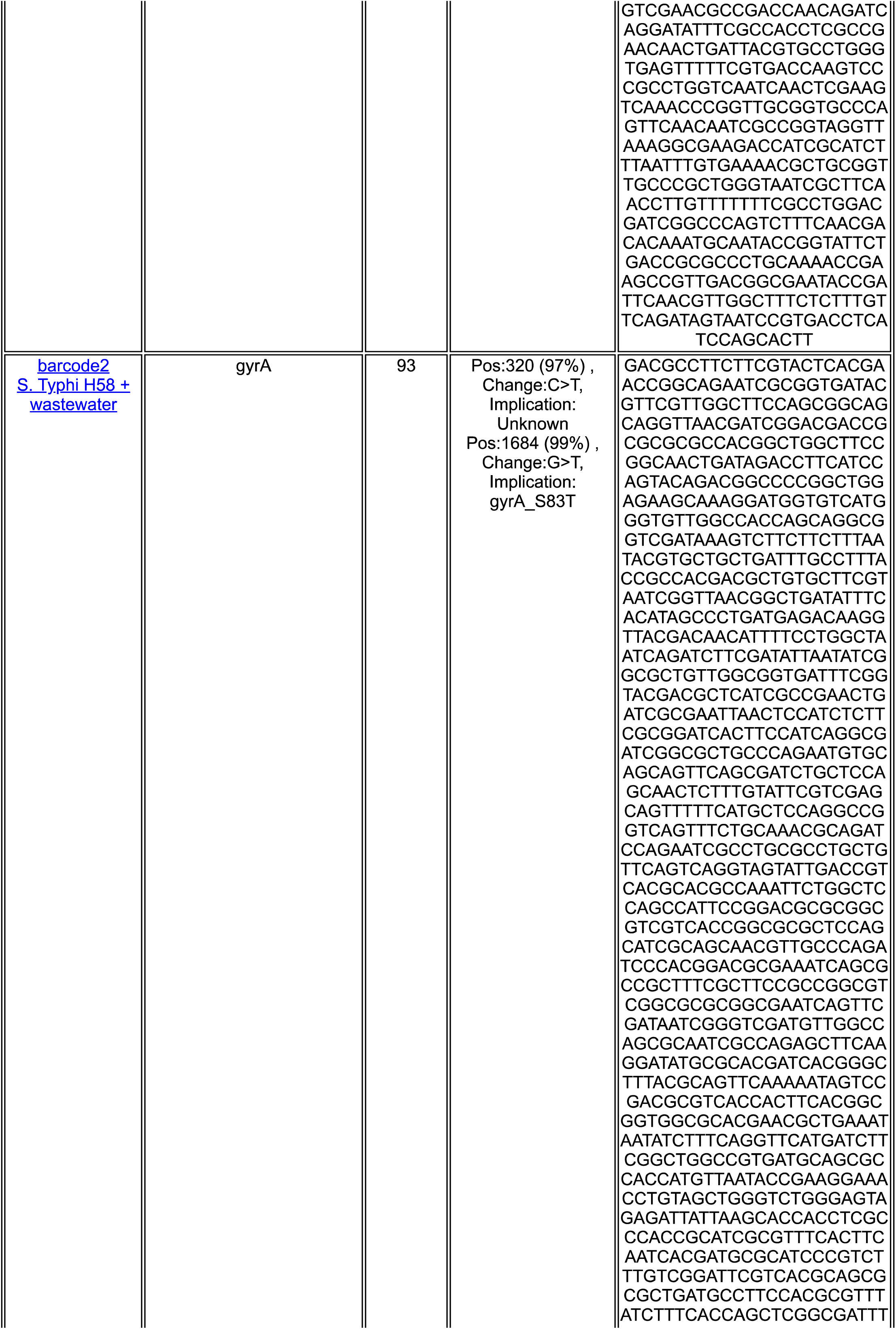
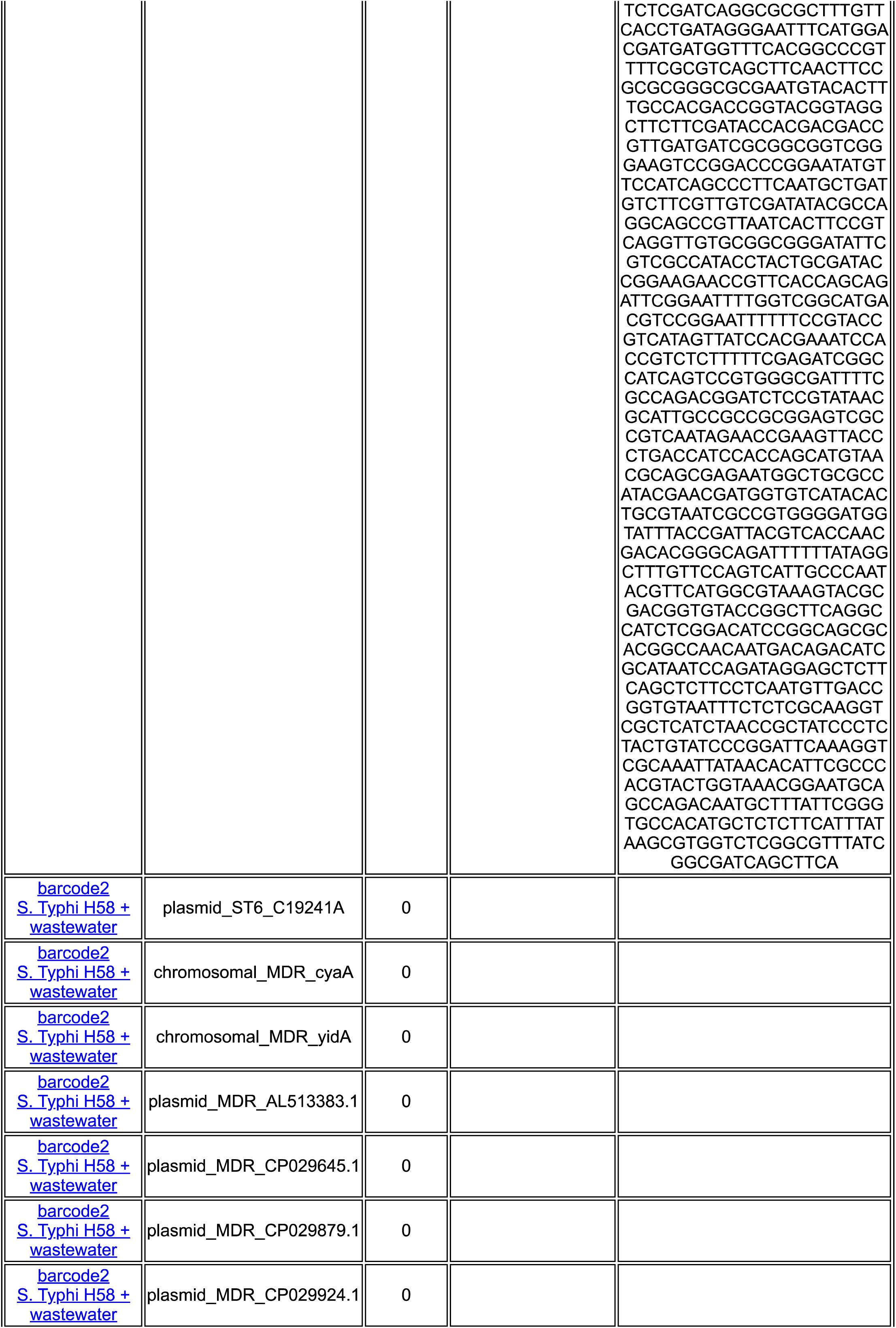
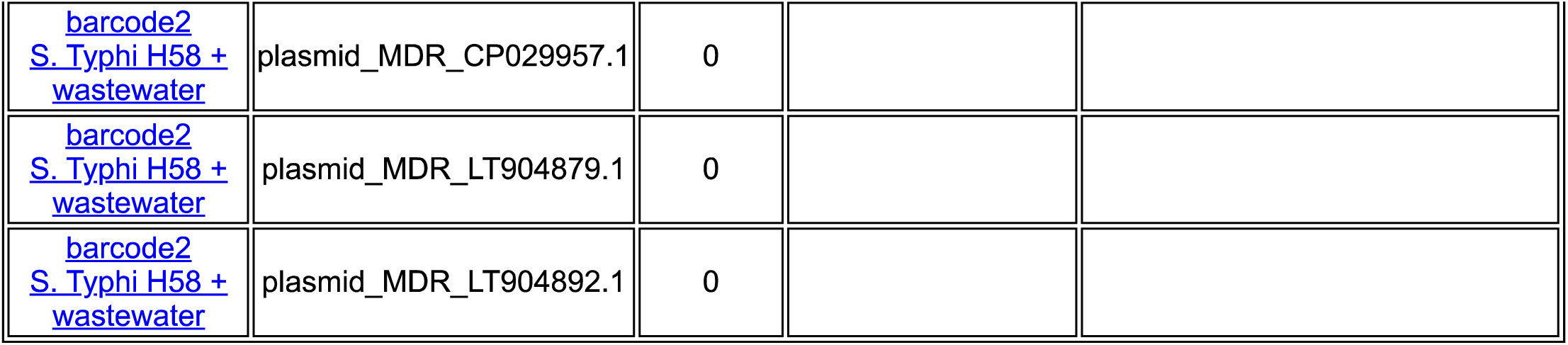

### Model Signatures

**Back to Summary**

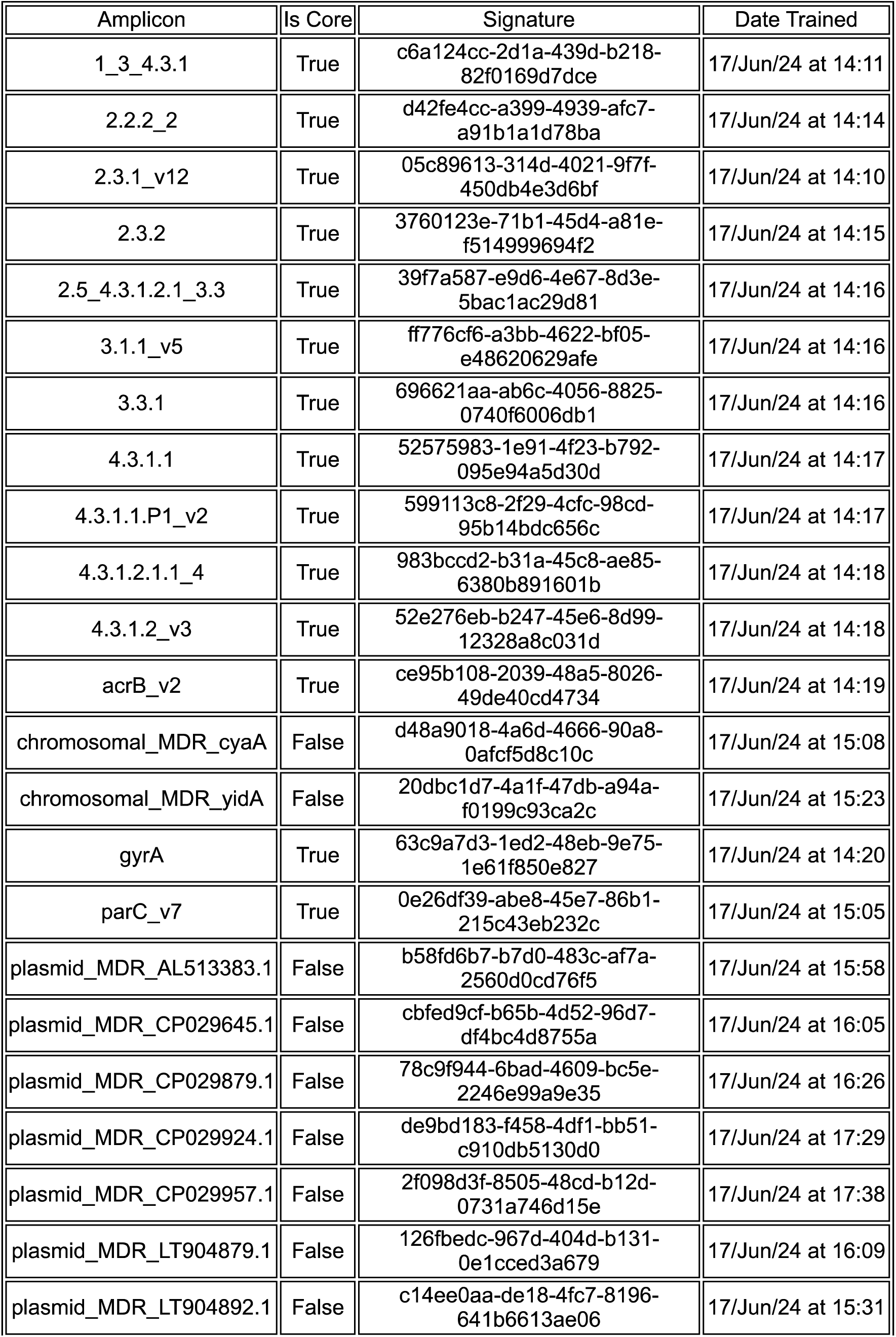
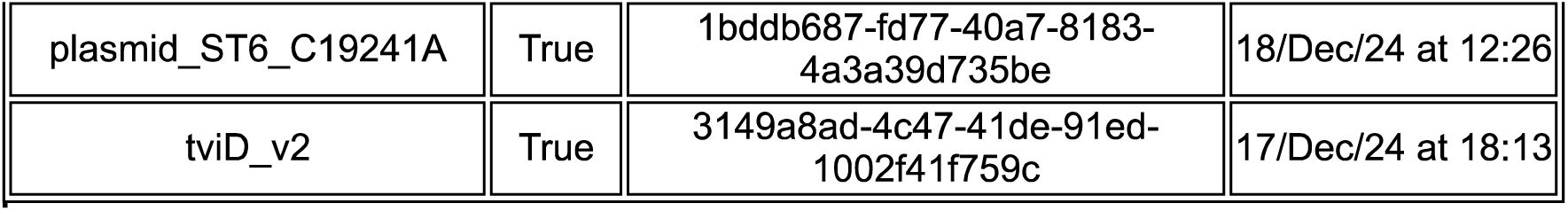

